## Supplementary Material for "Conditional prediction of consecutive tumor evolution using cancer progression models: What genotype comes next?"

2021-05-10 (Release: Rev: f614649)

### 1 Methods

#### 1.1 Transition probabilities from evolutionary simulations: supplementary details

Let us consider a matrix **P** where rows correspond to observable genotypes and columns to genotypes of the LOD. The element $(i, j)$ of the matrix, denoted as $p_{ij}$, represents the conditional probability that the genotype $j$ (with $n + 1$ mutations) is an element of the LOD given the genotype $i$ (with $n$ mutations) has been observed. Naturally, in the absence of back mutations and when mutations accumulate one by one, $p_{ij}$ is defined only if $i$ and $j$ are exactly one mutation away from each other. Given how our question is formulated, all the mutations in $i$ need to also be present in $j$ only in SSWM, when $i$ would necessarily be the parent of $j$. But in a more general scenario (e.g., when SSWM does not hold) it is possible that a genotype $i$ with $n$ mutations is observed despite it not belonging to the LOD. Therefore the genotype $j$ (element of the LOD and with $n + 1$ mutations) need not be a direct descendant of $i$. The only condition for $p_{ij}$ to be defined is that $j$ has exactly one more mutation than $i$. (Thus, the number of entries in $p_{i,j}$ that could be non-zero is $\binom{N}{n+1}$, where $N$ is the total number of loci). We built a transition matrix **P** for every fitness landscape that we generated. To obtain the elements $p_{ij}$ of each simulation, we ran through all 20000 evolutionary processes that were simulated in the corresponding landscape. In each case, we first listed all observable genotypes (i.e. those that represented the majority of the population at some point during the simulated evolutionary process). Then we extracted the LOD of every process tracking back the ancestors of the fixated genotype (see definition and details in [18]; using implementation in [5]). We identified the genotypes in the LOD with one mutation more than the observable ones, and added 1 to the corresponding entries of **P**. Finally, we row-normalized **P** so that $\sum_j p_{ij} = 1$. All other elements (i.e., those that do not satisfy $n_{mut}(j) = n_{mut}(i) + 1$) were set to 0. Each element $p_{ij}$ can thus be interpreted as the fraction of the times that $j$ was in the LOD given that an observation of $i$ had been made.

##### 1.1.1 Special cases

When the end of the evolutionary process is reached there are no further transitions to genotypes of the LOD. Thus, matrix **P**, which provides the true probability distributions for the question *what genotype of the LOD has* n + 1 *mutations iven that a genotype with n mutations has been observed?* needs to add to additional states that correspond to two distinct special cases. When the observed genotype is the final, fixated one, we have simply reached the end of the evolutionary process; we handle this by adding to the **P** an "end" situation (denoted as $p_{i,end}$). If SSWM conditions are not satisfied, however, observable genotypes need not be a part of the LOD so that there could be observable genotypes with more mutated loci than, or as many mutated loci as, the final genotype that gets fixated. In other words, it is possible that a clone with $m \geq n$ mutations establishes in the tumor for a period of time even if eventually another clone with $n$ mutations is the one that fixates. We handle this by adding to the **P** a "none" situation ($p_{i,none}$). Downstream analyses are performed including these two probabilities ($p_{i,end}$, $p_{i,none}$) (and row normalization was actually carried out after including *end* and *none*).

#### 1.2 Transition probabilities from CPMs: supplementary details

For MHN transition probabilities between genotypes for both the competing exponentials and time-discretized methods, were computed with code we wrote, available in the repository (functions `do_MHN` and `trans_rate_to_trans_mat`). The expressions to obtain the rates of the Markov process parameterized by the MHN are those given by [14] (see equation 2 and Fig. 2). We then obtained the probabilities of transition given a transition using the same competing exponentials approach as for CBN and MCCBN (see below). Time-discretized transition matrices were obtained as explained in the main manuscript.

For CBN, MCCBN, OT, CAPRESE, and CAPRI we wrote code to obtain the fitness graphs from the DAGs of restrictions: the fitness graph show which genotypes can descend from which parent genotype (but see below).

For CAPRESE and CAPRI (both CAPRI_AIC and CAPRI_BIC) this is all we used, since for these methods it is not possible to obtain probabilities of transitions. Even if we can obtain conditional probabilities of events (using functions such as `TRONCO::as.conditional.probs`) and the conditional probability tables (with functions such as `TRONCO:::as.bnlnearn.network`) there seems to be no mechanism under CAPRI/CAPRESE to obtain the conditional probabilities of genotypes implied by the models. As explained in detail in section "CAPRI, CAPRESE, and probabilities of paths of tumor progression" in S4_Text Supplementary file of [6], what "CAPRI[CAPRESE] is doing is fitting a Bayesian Network to the observational data, with the DAG[tree] built so that arrows respect thetemporal priority and probability raising restrictions. But the CPTs themselves are not estimated parameters of a model that could be mapped into probabilities of paths. The CPTs of CAPRI[CAPRESE] seem to be the conditional probabilities of observing what we observe under the DAG[tree] and they incorporate errors (model errors and noise in the data), and can contain non-zero entries for child nodes when their parents are absent." Thus, obtaining conditional probabilities of transition between genotypes seems impossible. Therefore, for CAPRESE, CAPRI_AIC and CAPRI_BIC, all transition from a genotype to its possible descendants under the DAG of restrictions were made equiprobable. Note that this is different from [6]: in [6] paths were set equiprobable; here it is transitions between genotypes that are set as equiprobable.

In that context, it must be noted that, strictly, the DAGs of CAPRI (and CAPRESE) do not encode deterministic dependencies: genotypes that do not fulfil the restrictions of the DAGs can still be possible. The edges in the DAGs specify the most likely mutational paths ("most common evolutionary trajectories" in [2]), but other mutational paths can take place. This can be confirmed by noting that the conditional probability tables for a gene when its ancestor dependencies are not satisfied can be non-zero. Moreover, this is the reason why CAPRI can present as different two DAGs, say DAG1 and DAG2, where DAG1 is the transitive reduction of DAG2: if the DAGs encoded deterministic dependencies both models would be identical (see further details and examples in section "CAPRI, CAPRESE, and probabilities of paths of tumor progression" in S4_Text Supplementary file of [6]). Thus, given that with CAPRI and CAPRESE it is not possible to obtain probabilities of transition between genotypes from the CPTs, we have opted to set as "possible transitions" those that are possible according to the DAGs and as not possible transitions those that are not possible according to the DAG, as the DAGs would seem to encode the relevant or important or most common trajectories according to CAPRI/CAPRESE.

For CBN and MCCBN we obtained probabilities of transition, conditioned in a transition, similarly to [11, 13]. The steps were (see also p. i729 of [13] and examples in section "Computing probabilities of paths" in S4_Text Supplementary file of [6]):

1. Obtain the set of genotypes that can exist under the poset.
2. Obtain the transition rate matrix between genotypes from the $\lambda$s (e.g., what is shown in matrix S in Montazeri et al. [13]). As explained in Montazeri et al., "the non-zero off-diagonal elements of the transition matrix are the transition rates from each genotype to its successive genotypes in the genotype lattice, also shown in Figure 1(b)." See also legend of Figure 1: "(b). Directed transition rates among neighboring genotypes are shown on the edges of the lattice".
3. Set the diagonal of the previous matrix to 0 and for each row of the transition rate matrix, divide by $\sum \lambda$. Now the entries are probabilities of transition to each descendant genotype given a transition using the usual competing exponentials approach.

For OT, we used `ot.fit$parent$est.weight` to obtain the probabilities of transition to each descendant genotype; if the OT fit cannot return an error estimate, that operation fails and in such a case we used the `ot.fit$parent$obs.weight` component. These probabilities are not really returned by the model, since the OTs used are untimed oncogenetic trees [4, 16].

For CBN, MCCBN, and OT, we also obtained equiprobable transitions and these are methods OT_uw, CBN_uw, and MCCBN_uw (where "uw" stands for "unweighted"). Equiprobable transitions are obtained, as for CAPRESE and CAPRI, from the fitness graphs.

For CBN and MCCBN time-discretized transition matrices for were obtained as explained in the main manuscript from the transition rate matrices. Note that time-discretized transition matrices are not available for OT as OT does not provide rates.

#### 1.3 Mismatches in genotype observability

In the simulations, genotypes that represent the majority of the tumor for a period of time during its progression are observable. On the other hand, it is straightforward to derive that for a CPM observable genotypes are those that have a probability greater than zero of being visited. In other words, genotype $j$ is considered observable if $\hat{p}_{ij} > 0$ for at least some $i$, i.e. if it is possible to transition to $j$ from at least one previous state. If a genotype $j$ is not observable according to a CPM, all $\hat{p}_{jk}$ is undefined for all $k$ (no prediction can be made regarding what comes after $j$, since $j$ is not even observable). In certain situations, a genotype can be observable in the simulations but deemed otherwise by the CPM or vice-versa. How do we compare the true probabilities ($p_{ij}$) with the estimations provided by the CPM ($\hat{p}_{ij}$) in these cases?

As summarized in the table below, if the genotype under consideration ($i$) is not observable in the simulations we simply exclude it from further analyses: since it is not going to be observed in practice, it is irrelevant what the CPM predicts. Else, if the genotype is observable according to both simulations and CPM, we calculate the Jensen-Shannon distance normally as explained earlier. If it is observable in the simulations but not deemed so by the CPM, since no prediction is provided we use that of the null model instead ($\mathbf{p}_i^0$): under this simple null model, all transitions from genotype $i$ to all genotypes with $n+1$ mutations plus the *end* and *none* states (see section Special cases) are considered equiprobable (note that MHN and MHN_td provide predictions for all possible genotypes —i.e., all genotypes are observable under MHN and MHN_td). The following table shows all the possible cases:

| | | Simulations | |
|---|---|---|---|
| | | $i$ observable | $i$ not observable |
| Predictions | $i$ observable | $JS(\mathbf{p}_i, \hat{\mathbf{p}}_i)$ | – |
| | $i$ not observable | $JS(\mathbf{p}_i, \mathbf{p}_i^0)$ | – |

When we measure the agreement of the predictions of two methods (say, CBN vs. MHN or CBN vs. CBN_td) we use a similar procedure. Let the predictions of methods 1 and 2 be called $\hat{\mathbf{p}}_i^1$ and $\hat{\mathbf{p}}_i^2$. Comparisons are performed as shown in the next table:

| | | Predictions method 2 | |
|---|---|---|---|
| | | $i$ observable | $i$ not observable |
| Predictions method 1 | $i$ observable | $JS(\hat{\mathbf{p}}_i^1, \hat{\mathbf{p}}_i^2)$ | $JS(\hat{\mathbf{p}}_i^1, \hat{\mathbf{p}}_i^0)$ |
| | $i$ not observable | $JS(\hat{\mathbf{p}}_i^0, \hat{\mathbf{p}}_i^2)$ | $JS(\hat{\mathbf{p}}_i^0, \hat{\mathbf{p}}_i^0)$ |

Note that the second row, second column, $JS(\hat{\mathbf{p}}_i^0, \hat{\mathbf{p}}_i^0)$, is necessarily 0 since we are comparing the same predictions of the null model.

#### 1.4 Quantification of similarity between predicted and true transition probabilities: supplementary details

Consider the vector of true probabilities $\mathbf{p}_i = (p_{ia}, p_{ib}, \cdots, p_{iz})$, where $a, b, \cdots, z$ are all genotypes with one mutation more than $i$: this is nothing but the $i$-th row of the matrix $\mathbf{P}$ (excluding the elements that do not satisfy $n_{mut}(j) = n_{mut}(i) + 1$). Consider also the analogous vector of predictions from a CPM $\hat{\mathbf{p}}_i$. These were computed as specified in sections Transition probabilities from evolutionary simulations: supplementary details, Transition probabilities from CPMs: supplementary details, and Mismatches in genotype observability.

We measured the similarity between these two vectors using the Jensen-Shannon distance ($JS$) [3], the square root of the Jensen-Shannon divergence [12]. The Jensen-Shannon divergence is a symmetrized Kullback-Leibler divergence between two probability distributions that is 0 when the two distributions are identical and reaches its maximum value of 1 if the two distributions do not overlap (we used log of base 2); it is defined even if the two distributions do not have the same sample space (i.e., even if $p_i \neq 0$ and $q_i = 0$ or viceversa). (And the square root of the Jensen-Shannon divergence is a metric between probability distributions [3])

#### 1.5 CPMs: software and usage details

For MHN [14], we used the software available from https://github.com/RudiSchill/MHN (downloaded in January 2020; as of 2020-11-18, that repository was last updated on 2018-08-16). For the tunning parameter $\lambda$ we used $1/|D|$ (see p. 244 of [14]) for the biological data. For the simulation data, since the true size of the data is known to us, but not the methods (as some data sets might have fewer features than the true ones), and to make analysis more easily reproducible and comparable, we used $\lambda = 0.01$ (same value as used in the original bioRxiv version of [14]).

For CBN, MCCBN, OT, CAPRESE, and CAPRI we have used the output of the analyses reported in [6], which provide full details on software and parameters for software in their S4_Text file. We summarize and update them here.

For CBN, we used version 0.1.04b from March 2016, and still current as of November 2020, downloaded from https://www.bsse.ethz.ch/cbg/software/ct-cbn.html. In the software repository we include the code we used (file ct-cbn-0.1.04b-with-rdu-bug-fix-write-lambda.tar.gz), which we have modified from the original; the changes include a bug fix for an unjustified stopping when (loglik_new < loglik; around lines 1434 of ct-cbn.h) and forcing ct-cbn to output a lambda_i, for a lambda from final iteration (the additions are around line 735 of ct-cbn.h). We wrote a wrapper to call CBN from R, and we used the default settings for temp ($-T = 1$) and steps ($-N =$ number of nodes$^2$) —tough we ensure a minimum of 25 steps are used, even if number of nodes is less than five; the simulated annealing search started for the best poset from an initial poset built using OT [17], as preliminary runs suggested this initial poset is as good as, or better than, the default linear poset in [10]. The parameters for the (exponentially distributed) waiting time until the occurrence of a mutation given its restrictions are satisfied ($\lambda$s) were obtained doing an additional run on the fitted model from the previous step, as in [10].

MCCBN was run using version 1.1.9 of the `mccbn` package, downloaded from github (https://github.com/cbg-ethz/MC-CBN on January 2018, which had been last updated on 2017-03-24).

OT was run using version 0.3.3 of the Oncotree package [17], still current as of November 2020.

CAPRI and CAPRESE were run using version 2.11.0 of the TRONCO BioConductor package, downloaded from the official BioConductor site. All options were left at the recommended defaults (e.g., 100 bootstrap samples for the estimation of the selective advantage scores with p-value of 0.05, and heuristic search using Hill Climbing).

#### 1.6 Simulated and cancer data sets

All simulated data sets were obtained from [6]. Before analyses, data from simulations were preprocessed as detailed in section "Preprocessing of data for CPMs" from Supplementary Material S4_Text of [6]. Briefly, all genes that were absent in all samples were removed (no inference can be made for these, since they are not observed); if two or more genes had identical observations over all individuals all except one were removed since these are indistinguishable events; if one or more genes were mutated in all samples one case (pseudosample) with no mutations (i.e., "wild-type") as added to the data set to allow us to use the exact same data for all methods without decreasing the dimensionality of the data set.

For the cancer data sets, In addition to the data sets in [6], we have used three CGH data sets previously used in [9, 14] obtained from the Progenetix database [1]. These are `Breast-CGH`, from 871 breast cancer patients with 10 CGH alterations, `Renal-CGH`, from 251 renal cell carcinoma with 12 CGH alterations, and the `Colon-CGH`, from 570 colorectal cancers with 11 CGH alterations. The data sets were obtained from the Supplementary data in [14].

#### 1.7 Linear mixed-effects model trees

##### 1.7.1 Linear mixed-effects model trees: brief description of procedure

To examine the relevance of the different factors on the performance of methods we used linear mixed-effects model trees. Linear mixed-effects model trees are an extension of recursive partitioning (or tree-based) methods. With recursive partitioning approaches, such as regression trees, instead of fitting a linear model where the dependent variable is expressed as a function of the predictor variables (possibly including interactions), observations are first split repeatedly according to the predictor variables, which play the role of partitioning variables, so that the dependent variable becomes more homogeneous within each node (i.e., so that we can find better-fitting models in the leaves —terminal nodes–, or local models) [7, 8]. Tree-based methods are especially interesting for exploratory work with many potential predictor variables and when interactions between predictor variables can be present, since they both automatically account for interactions (including high-order interactions) [8, 15] and they can facilitate interpretation by providing simpler models [7, 8]. Linear mixed-effects model trees extend tree-based methods by allowing us to take into account the dependency between observations due to shared random effects [7, 8, 15]. The linear mixed-effects model trees used here contain both global and local parts [8]. The global model uses all observation to fit the random-effects. The local model, in our case (since we are fitting models with constant fits in the leaves of the tree), only has an intercept, the estimated intercept for that leave or terminal node (recall that observations are partitioned over the tree with respect to the partitioning variables).

##### 1.7.2 Linear mixed-effects model trees: dependent and partitioning variables

The response variable, the minimal JS, was square root transformed before analysis as these resulted in closer agreement between the median and mean of the terminal nodes than the variable without transformation (data were asymmetrical).

Genotypes were weighted by their true proportion in samples (variable sampledProp). Weights were rescaled so the the sum of weights was equal to the total number of observations (number of rows of the data sets) (i.e., $weight = sampledProp \; (total\_number\_observations / \sum_i sampledProp_i)$. This weighting does not give more weight to data sets with larger sample size as, within a data set, $\sum_i sampledProp_i$ is 1. Note, though, that the weights were computed after removing the last genotype (and without rescaling the within-data set sampledProp); thus, genotypes from data sets where the last genotype was very common (i.e., data sets where the last genotype had larger sampledProp) are down-weighted. Other weighing schemes (with rescaling after removing the final genotype, for instance) could be devised. The consequences for analyses should be minor, though, since more than 90% of the data sets had a sum of sampledProp, after removing the last genotype, of 0.9 or larger.

All variables were treated as numerical variables except "typeLandscape", "detect" (detection regime), "nMut" (number of mutations of genotype), and "sample_size". "nMut" and "sample_size", even if they are integer-valued variables were fitted as categorical variables to allow for completely flexible splits, such as splits that leave non-contiguous number of mutations on the same side of the split (see, for example, node 3 in Figure **S1**, where observations with 0 and 8 mutations are on the right child node).

Type of fitness landscape, $\gamma$ (gamma), fraction of pairs of loci with reciprocal sign epistasis (epistRSign) and number of observed peaks in the fitness landscape (numObservedPeaks) were all used in the models, even when, as shown in Fig. **S4**, the RMF fitness landscape can be perfectly separated from the other two fitness landscapes by either number of observed peaks (numObservedPeaks) or fraction of reciprocal sign epistasis, and the local maxima and representable fitness landscapes can be almost perfectly separated by fraction of reciprocal sign epistasis. This was done on purpose to try to break down the contribution of fitness landscape into simpler factors.

##### 1.7.3 Linear mixed-effects model trees: fitting, parameters, pruning

Models were fitted using the R package "glmertree" [8].

The Bonferroni-corrected significance level for node-splitting was set at 0.01: argument `alpha = 0.01`. We specified a minimal size in leaves (or terminal nodes) of 1%: argument `minsize = 0.01 * nrow(dataset)`. Models were fit with five different optimizers, to check possible warnings during convergence. None of the fits used had warnings and were identical to model fits with other optimizers that also gave no warnings.

As the returned trees had over 50 leaves in all cases, to allow for interpretability we pruned the resulting tree by recursively merging (starting from the leaves) all children node with a fitted minimal JS > 0.0677 (which corresponds to the 20% best JS —see Table **S4**). In addition, we merged all children with a fitted minimal JS that differed by less than 0.0677/4 (so as to collapse good performing nodes with minor differences).

##### 1.7.4 Linear mixed-effects model trees: figure details

Figures below and in the main paper show the models after the pruning procedure and the displays of the internal nodes also give information about how the variables were treated. The internal nodes display histograms for categorical variables (such as "typeLandscape"), where the Y-axis is labelled "Freq" (frequency). Numerical variables show histograms (with Y-axis labelled "count") for integer-valued variables ("fitnessRank" "numObservedPeaks") and density plots for real-valued variables.

### 2 Data & code availability

Code for the analyses in this article is available at https://github.com/rdiaz02/what_genotype_next.

### 3 Supplementary results

#### 3.1 Minimal JS for combinations of number of genes, sample size, and detection region

| Number of genes | Sample size (sample_size) | Detection regime (detect) | Mean | Median |
|---|---|---|---|---|
| 7 | 50 | large | 0.3749 | 0.2668 |
| 7 | 200 | large | 0.2847 | 0.1878 |
| 7 | 4000 | large | 0.2175 | 0.1425 |
| 7 | 50 | small | 0.3256 | 0.2473 |
| 7 | 200 | small | 0.2883 | 0.2199 |
| 7 | 4000 | small | 0.2736 | 0.2108 |
| 7 | 50 | uniform | 0.2603 | 0.1912 |
| 7 | 200 | uniform | 0.2269 | 0.1655 |
| 7 | 4000 | uniform | 0.2154 | 0.1581 |
| 10 | 50 | large | 0.5432 | 0.5795 |
| 10 | 200 | large | 0.4293 | 0.3353 |
| 10 | 4000 | large | 0.2972 | 0.2215 |
| 10 | 50 | small | 0.4075 | 0.3351 |
| 10 | 200 | small | 0.3409 | 0.2874 |
| 10 | 4000 | small | 0.3229 | 0.2775 |
| 10 | 50 | uniform | 0.3237 | 0.2585 |
| 10 | 200 | uniform | 0.2713 | 0.2213 |
| 10 | 4000 | uniform | 0.2600 | 0.2138 |

**Table S1.** (Weighted) Mean and median minimal JS for all combinations of number of genes, sample size, and detection region. Weights are proportional to the true proportion in samples (variable sampledProp). The minimal JS is computed as the minimum JS over all 13 methods.

For the above table as well as tables **S2** and **S3** note that, as explained in section 1.7, this weighting does not give more weight to data sets with larger sample size as, within a data set, $\sum_i sampledProp_i$ is 1. See further details in that section.

#### 3.2 Minimal JS for combinations of number of genes, sample size, detection region, and fitness landscape

| | | | Fitness landscape (typeLandscape) | | |
|---|---|---|---|---|---|
| Number of genes | Sample size (sample_size) | Detection regime (detect) | Represent. | Local maxima | RMF |
| 7 | 50 | large | 0.448 | 0.366 | 0.315 |
| | | small | 0.263 | 0.291 | 0.425 |
| | | uniform | 0.205 | 0.249 | 0.320 |
| | 200 | large | 0.278 | 0.279 | 0.296 |
| | | small | 0.209 | 0.253 | 0.404 |
| | | uniform | 0.165 | 0.214 | 0.293 |
| | 4000 | large | 0.132 | 0.227 | 0.288 |
| | | small | 0.185 | 0.241 | 0.395 |
| | | uniform | 0.151 | 0.202 | 0.284 |
| 10 | 50 | large | 0.702 | 0.575 | 0.361 |
| | | small | 0.375 | 0.367 | 0.482 |
| | | uniform | 0.293 | 0.305 | 0.370 |
| | 200 | large | 0.523 | 0.433 | 0.337 |
| | | small | 0.266 | 0.301 | 0.456 |
| | | uniform | 0.211 | 0.255 | 0.340 |
| | 4000 | large | 0.267 | 0.293 | 0.330 |
| | | small | 0.234 | 0.283 | 0.451 |
| | | uniform | 0.196 | 0.243 | 0.334 |

**Table S2.** (Weighted) Mean minimal JS for all combinations of number of genes, sample size, detection region, and fitness landscape. Weights are proportional to the true proportion in samples (variable sampledProp). The minimal JS is computed as the minimum JS over all 13 methods.

| | | | Fitness landscape (typeLandscape) | | |
|---|---|---|---|---|---|
| Number of genes | Sample size (sample_size) | Detection regime (detect) | Represent. | Local maxima | RMF |
| 7 | 50 | large | 0.428 | 0.251 | 0.233 |
| | | small | 0.202 | 0.222 | 0.388 |
| | | uniform | 0.157 | 0.177 | 0.239 |
| | 200 | large | 0.150 | 0.191 | 0.220 |
| | | small | 0.174 | 0.200 | 0.372 |
| | | uniform | 0.133 | 0.153 | 0.215 |
| | 4000 | large | 0.079 | 0.160 | 0.219 |
| | | small | 0.160 | 0.193 | 0.359 |
| | | uniform | 0.124 | 0.149 | 0.209 |
| 10 | 50 | large | 0.826 | 0.609 | 0.298 |
| | | small | 0.291 | 0.294 | 0.462 |
| | | uniform | 0.241 | 0.235 | 0.302 |
| | 200 | large | 0.678 | 0.322 | 0.286 |
| | | small | 0.232 | 0.258 | 0.438 |
| | | uniform | 0.186 | 0.204 | 0.283 |
| | 4000 | large | 0.180 | 0.220 | 0.284 |
| | | small | 0.215 | 0.249 | 0.435 |
| | | uniform | 0.177 | 0.194 | 0.280 |

**Table S3.** (Weighted) Median minimal JS for all combinations of number of genes, sample size, detection region, and fitness landscape. Weights are proportional to the true proportion in samples (variable sampledProp). The minimal JS is computed as the minimum JS over all 13 methods.

##### 3.3 Minimal JS and JS for each method: quantiles

| | Minimal | MHN | CBN | MHN_td | CBN_td | MCCBN | MCCBN_td | CAPRI_AIC | CAPRI_BIC | CBN_uw | MCCBN_uw | CAPRESE | OT | OT_uw |
|---|---|---|---|---|---|---|---|---|---|---|---|---|---|---|
| 0 | 0.000 | 0.000 | 0.000 | 0.000 | 0.000 | 0.000 | 0.000 | 0.000 | 0.000 | 0.000 | 0.000 | 0.000 | 0.000 | 0.000 |
| 0.05 | 0.000 | 0.076 | 0.000 | 0.059 | 0.061 | 0.000 | 0.077 | 0.000 | 0.005 | 0.000 | 0.000 | 0.000 | 0.000 | 0.000 |
| 0.1 | 0.017 | 0.150 | 0.075 | 0.122 | 0.138 | 0.065 | 0.176 | 0.121 | 0.139 | 0.105 | 0.075 | 0.094 | 0.086 | 0.098 |
| 0.15 | 0.042 | 0.195 | 0.134 | 0.218 | 0.256 | 0.129 | 0.318 | 0.239 | 0.258 | 0.199 | 0.159 | 0.187 | 0.166 | 0.193 |
| 0.2 | 0.068 | 0.233 | 0.187 | 0.353 | 0.416 | 0.191 | 0.483 | 0.339 | 0.363 | 0.285 | 0.235 | 0.268 | 0.236 | 0.279 |
| 0.25 | 0.094 | 0.269 | 0.237 | 0.588 | 0.594 | 0.251 | 0.614 | 0.434 | 0.454 | 0.360 | 0.305 | 0.341 | 0.299 | 0.356 |
| 0.4 | 0.176 | 0.387 | 0.392 | 0.873 | 0.824 | 0.440 | 0.802 | 0.628 | 0.623 | 0.550 | 0.511 | 0.538 | 0.483 | 0.558 |
| 0.5 | 0.237 | 0.500 | 0.551 | 0.930 | 0.898 | 0.663 | 0.868 | 0.767 | 0.741 | 0.624 | 0.678 | 0.628 | 0.634 | 0.671 |
| 0.75 | 0.467 | 0.935 | 0.962 | 0.983 | 0.982 | 0.962 | 0.954 | 0.969 | 0.968 | 0.960 | 0.962 | 0.960 | 0.963 | 0.962 |
| 1 | 1.000 | 1.000 | 1.000 | 1.000 | 1.000 | 1.000 | 1.000 | 1.000 | 1.000 | 1.000 | 1.000 | 1.000 | 1.000 | 1.000 |

**Table S4.** (Weighted) Quantiles of minimal JS and JS for each method for the complete data set.

| | Minimal | MHN | CBN | MHN_td | CBN_td | MCCBN | MCCBN_td | CAPRI_AIC | CAPRI_BIC | CBN_uw | MCCBN_uw | CAPRESE | OT | OT_uw |
|---|---|---|---|---|---|---|---|---|---|---|---|---|---|---|
| 0 | 0.000 | 0.000 | 0.000 | 0.118 | 0.000 | 0.000 | 0.000 | 0.000 | 0.000 | 0.000 | 0.000 | 0.000 | 0.000 | 0.000 |
| 0.05 | 0.000 | 0.000 | 0.000 | 0.634 | 0.390 | 0.000 | 0.342 | 0.000 | 0.000 | 0.000 | 0.000 | 0.000 | 0.000 | 0.000 |
| 0.1 | 0.000 | 0.000 | 0.000 | 0.750 | 0.540 | 0.000 | 0.466 | 0.000 | 0.000 | 0.000 | 0.000 | 0.000 | 0.000 | 0.000 |
| 0.15 | 0.000 | 0.000 | 0.000 | 0.805 | 0.612 | 0.000 | 0.534 | 0.000 | 0.000 | 0.000 | 0.000 | 0.000 | 0.000 | 0.000 |
| 0.2 | 0.000 | 0.101 | 0.000 | 0.838 | 0.664 | 0.000 | 0.583 | 0.042 | 0.034 | 0.000 | 0.000 | 0.000 | 0.000 | 0.000 |
| 0.25 | 0.023 | 0.137 | 0.036 | 0.863 | 0.700 | 0.035 | 0.624 | 0.149 | 0.136 | 0.049 | 0.045 | 0.049 | 0.062 | 0.064 |
| 0.4 | 0.104 | 0.212 | 0.135 | 0.911 | 0.789 | 0.138 | 0.721 | 0.382 | 0.359 | 0.180 | 0.176 | 0.191 | 0.223 | 0.233 |
| 0.5 | 0.149 | 0.257 | 0.193 | 0.932 | 0.836 | 0.200 | 0.772 | 0.505 | 0.489 | 0.252 | 0.246 | 0.268 | 0.311 | 0.322 |
| 0.75 | 0.266 | 0.377 | 0.337 | 0.971 | 0.920 | 0.354 | 0.883 | 0.835 | 0.829 | 0.421 | 0.410 | 0.451 | 0.531 | 0.546 |
| 1 | 1.000 | 1.000 | 1.000 | 1.000 | 1.000 | 1.000 | 1.000 | 1.000 | 1.000 | 1.000 | 1.000 | 1.000 | 1.000 | 1.000 |

**Table S5.** (Weighted) Quantiles of minimal JS and JS for each method for representable fitness landscapes, sample size of 4000, and uniform detection regime.

| | Minimal | MHN | CBN | MHN_td | CBN_td | MCCBN | MCCBN_td | CAPRI_AIC | CAPRI_BIC | CBN_uw | MCCBN_uw | CAPRESE | OT | OT_uw |
|---|---|---|---|---|---|---|---|---|---|---|---|---|---|---|
| 0 | 0.000 | 0.000 | 0.000 | 0.000 | 0.000 | 0.000 | 0.000 | 0.000 | 0.000 | 0.000 | 0.000 | 0.000 | 0.000 | 0.000 |
| 0.05 | 0.000 | 0.132 | 0.036 | 0.032 | 0.045 | 0.000 | 0.047 | 0.006 | 0.019 | 0.062 | 0.000 | 0.006 | 0.002 | 0.001 |
| 0.1 | 0.015 | 0.178 | 0.093 | 0.059 | 0.085 | 0.062 | 0.095 | 0.136 | 0.147 | 0.193 | 0.078 | 0.112 | 0.084 | 0.106 |
| 0.15 | 0.030 | 0.217 | 0.140 | 0.094 | 0.124 | 0.123 | 0.144 | 0.252 | 0.270 | 0.305 | 0.161 | 0.203 | 0.163 | 0.199 |
| 0.2 | 0.045 | 0.249 | 0.182 | 0.139 | 0.177 | 0.180 | 0.208 | 0.374 | 0.389 | 0.390 | 0.239 | 0.290 | 0.233 | 0.291 |
| 0.25 | 0.064 | 0.287 | 0.230 | 0.235 | 0.255 | 0.238 | 0.281 | 0.473 | 0.490 | 0.452 | 0.310 | 0.374 | 0.304 | 0.379 |
| 0.4 | 0.123 | 0.399 | 0.361 | 0.677 | 0.573 | 0.393 | 0.546 | 0.678 | 0.675 | 0.558 | 0.511 | 0.554 | 0.487 | 0.557 |
| 0.5 | 0.171 | 0.509 | 0.492 | 0.828 | 0.708 | 0.555 | 0.665 | 0.860 | 0.859 | 0.607 | 0.603 | 0.619 | 0.620 | 0.648 |
| 0.75 | 0.329 | 0.980 | 0.981 | 0.949 | 0.900 | 0.978 | 0.856 | 0.979 | 0.980 | 0.981 | 0.978 | 0.980 | 0.981 | 0.981 |
| 1 | 1.000 | 1.000 | 1.000 | 1.000 | 1.000 | 1.000 | 1.000 | 1.000 | 1.000 | 1.000 | 1.000 | 1.000 | 1.000 | 1.000 |

**Table S6.** (Weighted) Quantiles of minimal JS and JS for each method for local maxima fitness landscapes, sample size of 4000, and uniform detection regime.

| | Minimal | MHN | CBN | MHN_td | CBN_td | MCCBN | MCCBN_td | CAPRI_AIC | CAPRI_BIC | CBN_uw | MCCBN_uw | CAPRESE | OT | OT_uw |
|---|---|---|---|---|---|---|---|---|---|---|---|---|---|---|
| 0 | 0.000 | 0.000 | 0.000 | 0.000 | 0.000 | 0.000 | 0.000 | 0.000 | 0.000 | 0.000 | 0.000 | 0.000 | 0.000 | 0.000 |
| 0.05 | 0.011 | 0.145 | 0.122 | 0.055 | 0.030 | 0.109 | 0.042 | 0.247 | 0.277 | 0.315 | 0.144 | 0.244 | 0.149 | 0.244 |
| 0.1 | 0.037 | 0.205 | 0.215 | 0.093 | 0.071 | 0.233 | 0.101 | 0.381 | 0.407 | 0.434 | 0.280 | 0.364 | 0.253 | 0.360 |
| 0.15 | 0.059 | 0.260 | 0.287 | 0.129 | 0.113 | 0.319 | 0.165 | 0.486 | 0.524 | 0.511 | 0.379 | 0.471 | 0.337 | 0.470 |
| 0.2 | 0.083 | 0.325 | 0.355 | 0.169 | 0.180 | 0.386 | 0.234 | 0.559 | 0.565 | 0.558 | 0.470 | 0.551 | 0.415 | 0.550 |
| 0.25 | 0.104 | 0.385 | 0.420 | 0.208 | 0.246 | 0.465 | 0.321 | 0.602 | 0.630 | 0.578 | 0.556 | 0.575 | 0.487 | 0.571 |
| 0.4 | 0.182 | 0.596 | 0.716 | 0.392 | 0.611 | 0.813 | 0.631 | 0.855 | 0.855 | 0.739 | 0.816 | 0.753 | 0.743 | 0.760 |
| 0.5 | 0.245 | 0.771 | 0.893 | 0.618 | 0.797 | 0.915 | 0.781 | 0.930 | 0.931 | 0.891 | 0.915 | 0.904 | 0.906 | 0.906 |
| 0.75 | 0.465 | 0.998 | 0.999 | 0.906 | 0.980 | 0.998 | 0.935 | 0.999 | 0.999 | 0.999 | 0.998 | 0.999 | 0.999 | 0.999 |
| 1 | 1.000 | 1.000 | 1.000 | 1.000 | 1.000 | 1.000 | 1.000 | 1.000 | 1.000 | 1.000 | 1.000 | 1.000 | 1.000 | 1.000 |

**Table S7.** (Weighted) Quantiles of minimal JS and JS for each method for RMF fitness landscapes, sample size of 4000, and uniform detection regime.

##### 3.4 Linear mixed-effects model trees

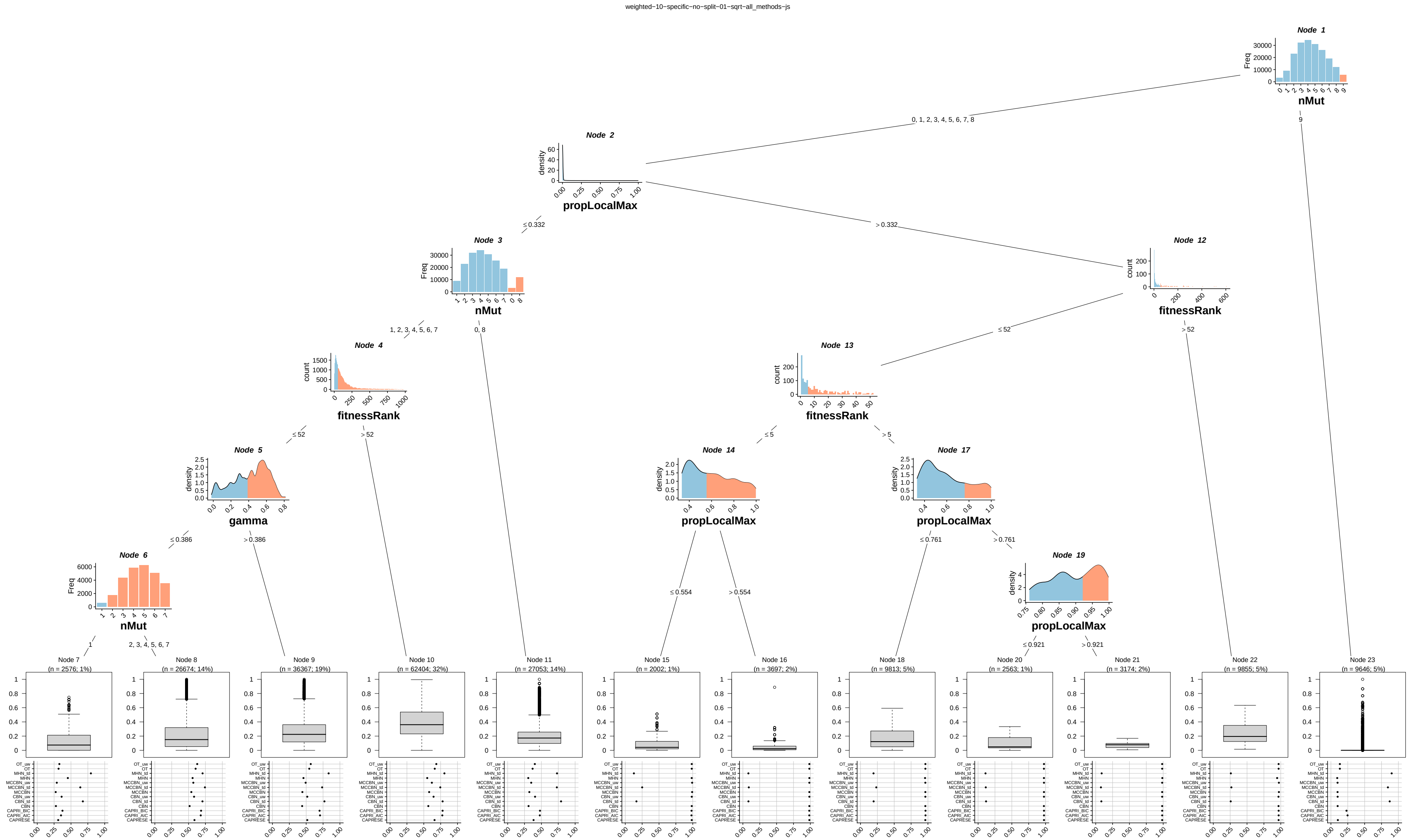

**Fig. S1. Linear mixed-effects model tree to data from fitness landscapes with 10 loci, sample size of 4000 and uniform sampling.** Internal nodes show the partitioning variables and the values that lead to each partition; light blue denotes the values or categories that lead down the left split and salmon the values or categories that lead down the right split (partitioning criteria are also shown in the edges leading down a node). Leaves or terminal nodes show (weighted) boxplots of the dependent variable for the observations that belong that fall in that leave and, as label, the name of the node and the sample size (sum of weighted observations and % of the total data set for the fit). The dot plots underneath the boxplots show the mean JS for all the methods for that leave.

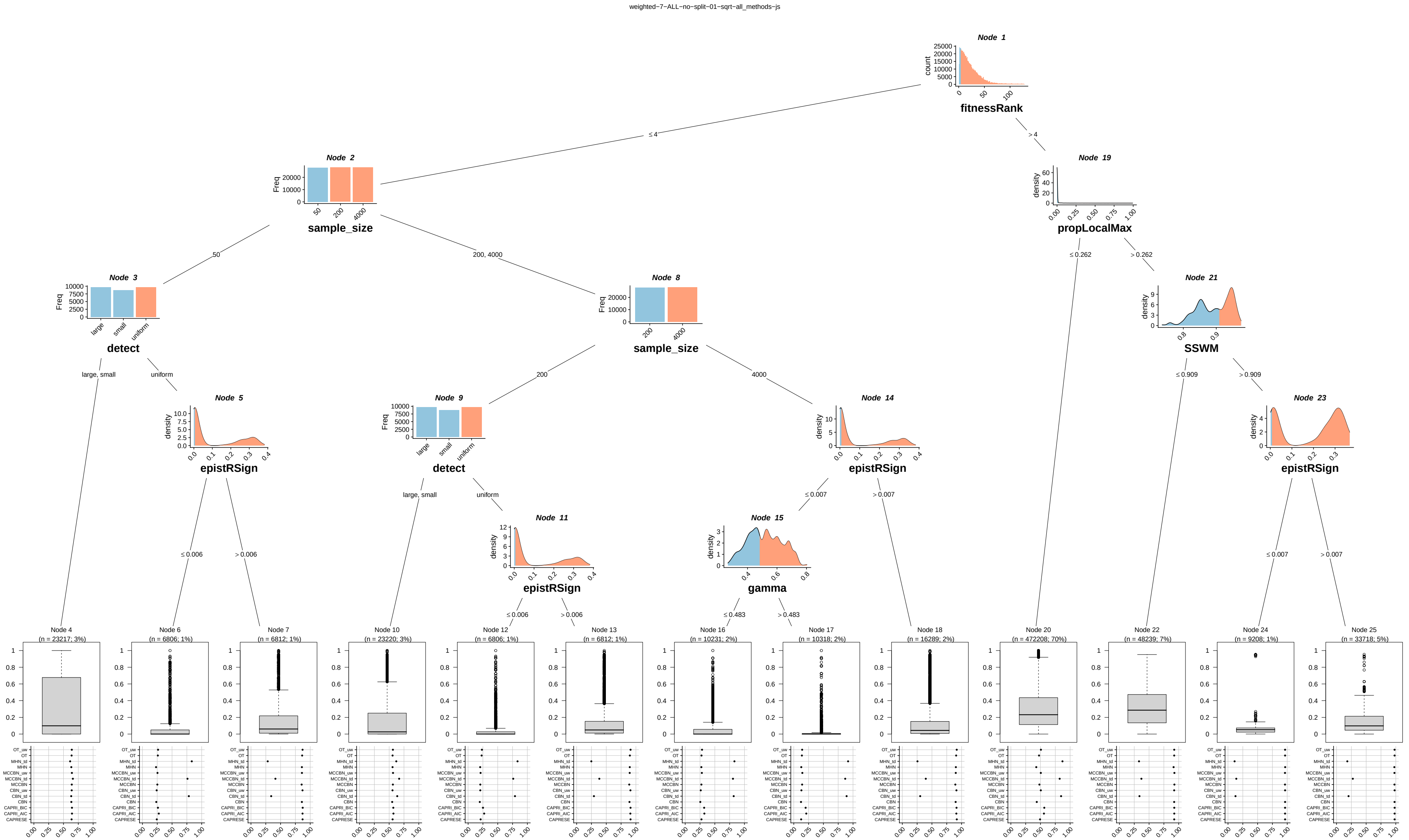

**Fig. S2. Linear mixed-effects model tree to all data from fitness landscapes with 7 loci (or genes).** Internal nodes show the partitioning variables and the values that lead to each partition; light blue denotes the values or categories that lead down the left split and salmon the values or categories that lead down the right split (partitioning criteria are also shown in the edges leading down a node). Leaves or terminal nodes show (weighted) boxplots of the dependent variable for the observations that belong that fall in that leave and, as label, the name of the node and the sample size (sum of weighted observations and % of the total data set for the fit). The dot plots underneath the boxplots show the mean JS for all the methods for that leave.

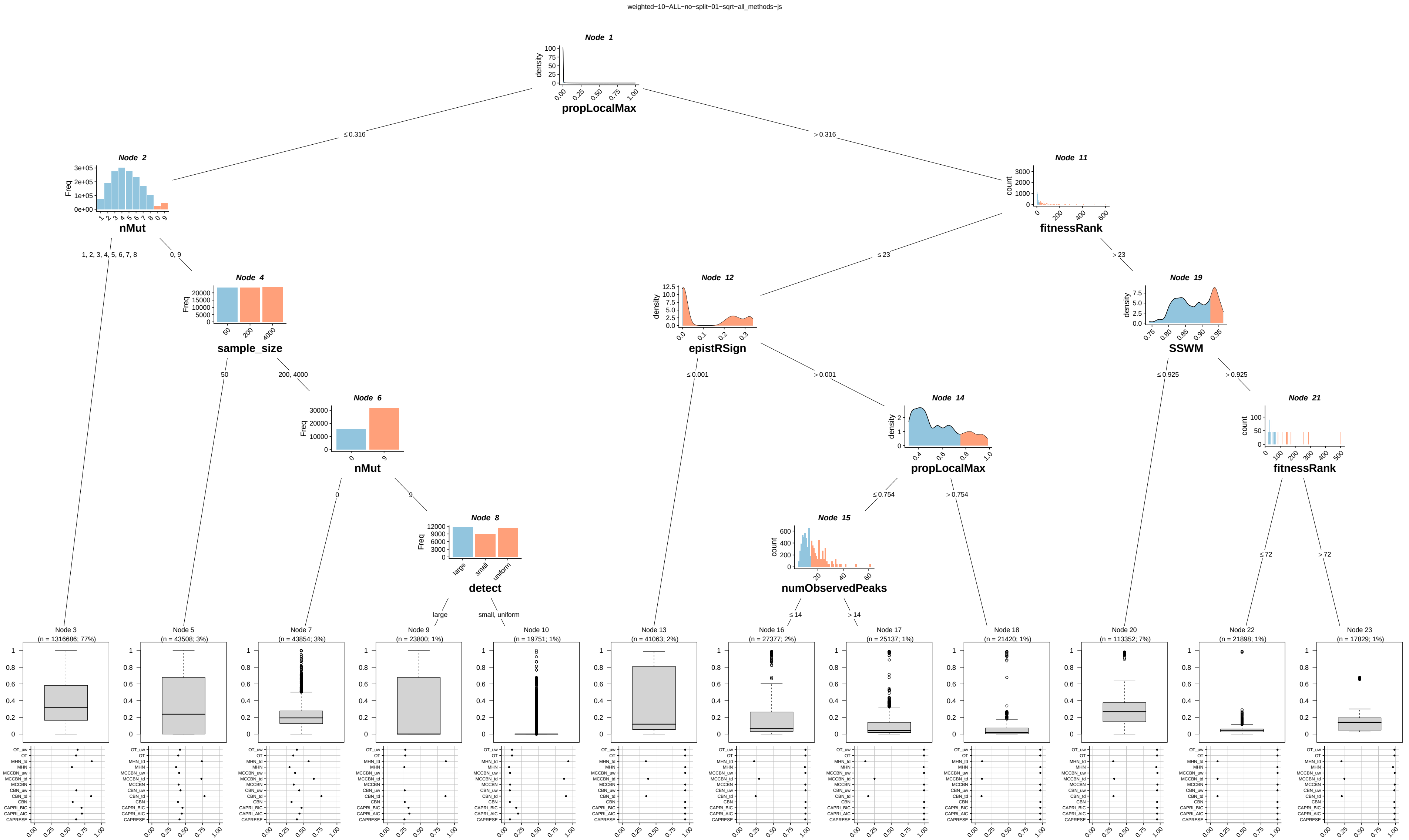

**Fig. S3. Linear mixed-effects model tree to all data from fitness landscapes with 10 loci (or genes).** Internal nodes show the partitioning variables and the values that lead to each partition; light blue denotes the values or categories that lead down the left split and salmon the values or categories that lead down the right split (partitioning criteria are also shown in the edges leading down a node). Leaves or terminal nodes show (weighted) boxplots of the dependent variable for the observations that belong that fall in that leave and, as label, the name of the node and the sample size (sum of weighted observations and % of the total data set for the fit). The dot plots underneath the boxplots show the mean JS for all the methods for that leave.

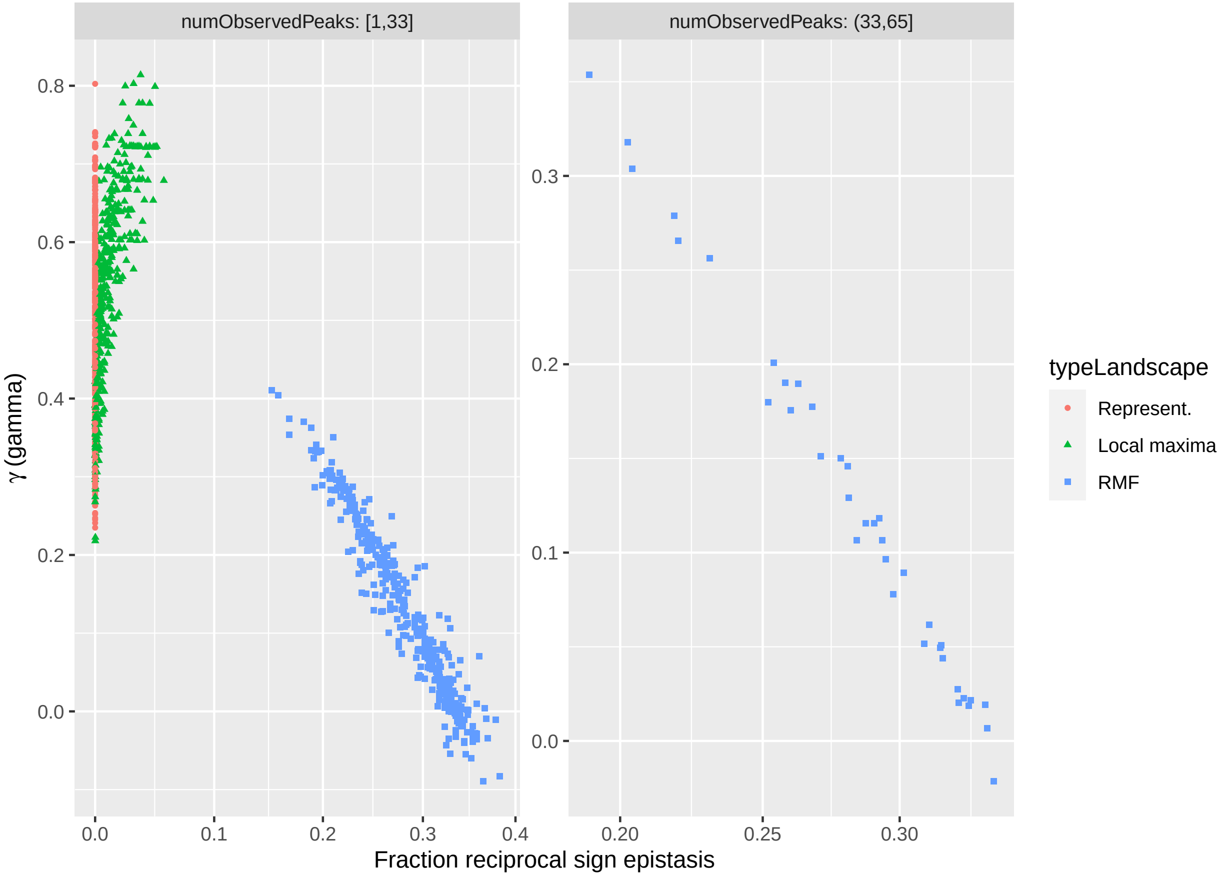

**Fig. S4. Relationship between $\gamma$, reciprocal sign epistasis, number of observed peaks, and fitness landscape.** The RMF fitness landscape can be perfectly separated from the other two fitness landscapes by either number of observed peaks (numObservedPeaks) or fraction of reciprocal sign epistasis. Number of observed peaks and fraction of reciprocal sign epistasis are constant (1 and 0, respectively) for the "Representable" fitness landscapes; these values are also seen in some of the Local maxima fitness landscapes (as even if the local maxima fitness landscapes do indeed have local maxima, that need not mean that the actual evolutionary process will, with probability 1, result in more than one maxima being visited); therefore, these two types of fitness landscapes cannot be perfectly separated using these three variables. Finally, notice how the relationship between $\gamma$ and reciprocal sign epistasis (epistRSign in previous figures) differs between the RMF and the local maxima fitness landscapes.

#### 3.5 Categorization of similarities between methods and performance

The following tables summarize scenarios of good performance. All tables present weighted sums and/or proportions. Weights are proportional to the true proportion in samples (variable sampledProp). Variables in tables have the following meaning:

**Good TD performance** True when the JS (from comparing predictions with the truth) of both CBN_td and MHN_td ≤ 0.0677. We exclude MCCBN_td here because of its slightly worse performance.

**Good CE performance** True when the JS (from comparing predictions with the truth) of both CBN and MHN ≤ 0.0677. We exclude MCCBN for consistency with the previous statistic.

Thus, the above two variables summarize the answer to the question "could we do a good job with one of the methods in TD (CBN_td, MHN_td) or one of the methods in the CE set?". It is an answer to "is good performance with any of the candidates (in one of the sets) possible?"

**Similar TD** True when the similarity of predictions (measured with JS) between CBN_td and MHN_td ≤ 0.0677.

**Similar CE** True when the similarity of predictions (measured with JS) between CBN and MHN ≤ 0.0677.

**Different CE-TD** We measured the similarity of predictions (measured with JS) between the CE and TD methods, defined as the average of the similarity between CBN and CBN_td and MHN and MHN_td. (MCCBN was not used as it showed slightly poorer performance). Precisely, this variable is true when $(1/2)\ (JS_{CBN,CBN\_td} + JS_{MHN,MHN\_td}) \geq 0.7$.

The above variables measure if the output of different methods or families of methods are similar.

Note that extending the "CE" to include CAPRESE and OT changes very little. For simplicity, and because of their slightly better performance, we focus here only on CBN, MCCBN, and MHN.

The tables below present straightforward conditional probability calculations for good performance given different patterns of similarity between and among methods. Note that these are calculations based on the observed data, and not rigorous estimates of classification accuracy, that would require cross-validation or bootstrap with possibly more sophisticated metrics. We show them here simply to illustrate upper bounds on performance.

| Different CE-TD | Similar TD | Good TD FALSE | Good TD TRUE |
|---|---|---|---|
| FALSE | FALSE | 0.185 | 0.000 |
| | TRUE | 0.117 | 0.002 |
| TRUE | FALSE | 0.048 | 0.000 |
| | TRUE | 0.611 | 0.037 |

**Table S8.** Cross-tabulation of Good TD, Similar TD, and Different CE-TD. Values shown are (weighted) proportion of cases (proportion computed over the sum of all cells). The cells with 0 are necessarily 0 by the definition of "Good TD".
$P(\text{Good TD}|\text{SimilarTD}) = 0.039/(0.037 + 0.002 + 0.611 + 0.117) = 0.0508$.
$P(\text{Good TD}|\text{Similar TD \& Different CE} - \text{TD}) = 0.037/(0.037 + 0.611) = 0.057$.
$P(\text{Good TD}) = 0.039$.

| Different CE-TD | Similar CE | Good CE FALSE | Good CE TRUE |
|---|---|---|---|
| FALSE | FALSE | 0.155 | 0.000 |
| | TRUE | 0.147 | 0.003 |
| TRUE | FALSE | 0.440 | 0.000 |
| | TRUE | 0.218 | 0.038 |

**Table S9.** Cross-tabulation of Good CE, Similar CE, and Different CE-TD. Values shown are (weighted) proportion of cases (proportion computed over the sum of all cells). The cells with 0 are necessarily 0 by the definition of "Good TD".
$P(\text{Good CE}|\text{SimilarCE}) = 0.041/(0.038 + 0.003 + 0.218 + 0.147) = 0.101$.
$P(\text{Good CE}|\text{Similar CE \& Different CE} - \text{TD}) = 0.038/(0.038 + 0.218) = 0.148$.
$P(\text{Good CE}) = 0.041$.

| | | | Good TD | |
|---|---|---|---|---|
| Different CE-TD | Similar TD | Similar CE | FALSE | TRUE |
| FALSE | FALSE | FALSE | 0.140 | 0.000 |
| | | TRUE | 0.045 | 0.000 |
| | TRUE | FALSE | 0.014 | 0.000 |
| | | TRUE | 0.103 | 0.002 |
| TRUE | FALSE | FALSE | 0.034 | 0.000 |
| | | TRUE | 0.014 | 0.000 |
| | TRUE | FALSE | 0.375 | 0.030 |
| | | TRUE | 0.236 | 0.006 |

**Table S10.** Cross-tabulation of Good TD, Similar CE, Similar TD, and Different CE-TD.
$P(\text{Good TD}|\text{Similar CE \& Similar TD \& Different CE} - \text{TD}) = 0.006/(0.006 + 0.236) = 0.025$.
$P(\text{Good TD}|\neg\text{Similar CE \& Similar TD \& Different CE} - \text{TD}) = 0.030/(0.030 + 0.375) = 0.074$.

| | | | Good CE | |
|---|---|---|---|---|
| Different CE-TD | Similar TD | Similar CE | FALSE | TRUE |
| FALSE | FALSE | FALSE | 0.140 | 0.000 |
| | | TRUE | 0.044 | 0.001 |
| | TRUE | FALSE | 0.014 | 0.000 |
| | | TRUE | 0.103 | 0.002 |
| TRUE | FALSE | FALSE | 0.034 | 0.000 |
| | | TRUE | 0.013 | 0.001 |
| | TRUE | FALSE | 0.405 | 0.000 |
| | | TRUE | 0.204 | 0.038 |

**Table S11.** Cross-tabulation of Good CE, Similar CE, Similar TD, and Different CE-TD.
$P(\text{Good CE}|\text{Similar CE \& Similar TD \& Different CE} - \text{TD}) = 0.038/(0.204 + 0.038) = 0.157$.
$P(\text{Good CE}|\text{Similar CE \& }\neg\text{Similar TD \& Different CE} - \text{TD}) = 0.001/(0.001 + 0.013) = 0.071$.

##### 3.6 Similarity of predictions for all genotypes for the biological data sets

The figures in this section show the similarity of predictions for all genotypes for all data sets. The TD vs. TD comparison is $JS_{CBN\_td,MHN\_td}$, the CE vs. CE comparisons is $JS_{CBN,MHN}$, and the CE vs. TD comparison are $JS_{CBN,CBN\_td}$, $JS_{MHN,MHN\_td}$. As in the rest of the figures, the genotype with all mutations, if observed, has been excluded in these figures. Notice that some of the values of 0 for CE vs. TD comparisons are due to CBN not producing a prediction for that genotype, which therefore leads to identical equiprobable predictions from both CBN and CBN_td (see details in section Mismatches in genotype observability).

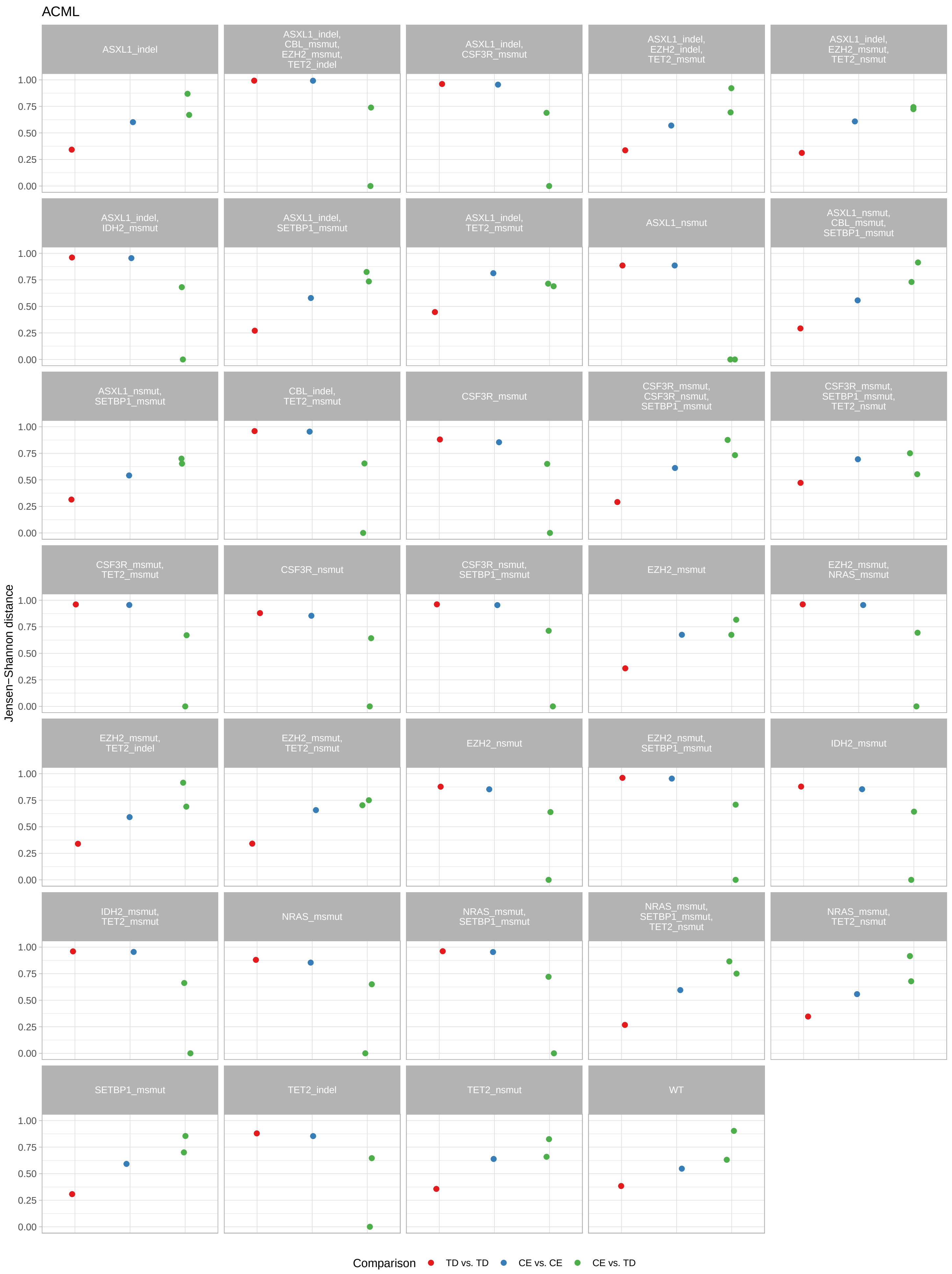

ACML
Jensen–Shannon distance
1.00
0.75
0.50
0.25
0.00
ASXL1_indel
ASXL1_indel, CBL_msmut, EZH2_msmut, TET2_indel
ASXL1_indel, CSF3R_msmut
ASXL1_indel, EZH2_indel, TET2_msmut
ASXL1_indel, EZH2_msmut, TET2_nsmut
ASXL1_indel, IDH2_msmut
ASXL1_indel, SETBP1_msmut
ASXL1_indel, TET2_msmut
ASXL1_nsmut
ASXL1_nsmut, CBL_msmut, SETBP1_msmut
ASXL1_nsmut, SETBP1_msmut
CBL_indel, TET2_msmut
CSF3R_msmut
CSF3R_msmut, CSF3R_nsmut, SETBP1_msmut
CSF3R_msmut, SETBP1_msmut, TET2_nsmut
CSF3R_msmut, TET2_msmut
CSF3R_nsmut
CSF3R_nsmut, SETBP1_msmut
EZH2_msmut
EZH2_msmut, NRAS_msmut
EZH2_msmut, TET2_indel
EZH2_msmut, TET2_nsmut
EZH2_nsmut
EZH2_nsmut, SETBP1_msmut
IDH2_msmut
IDH2_msmut, TET2_msmut
NRAS_msmut
NRAS_msmut, SETBP1_msmut
NRAS_msmut, SETBP1_msmut, TET2_nsmut
NRAS_msmut, TET2_nsmut
SETBP1_msmut
TET2_indel
TET2_nsmut
WT
Comparison
TD vs. TD
CE vs. CE
CE vs. TD

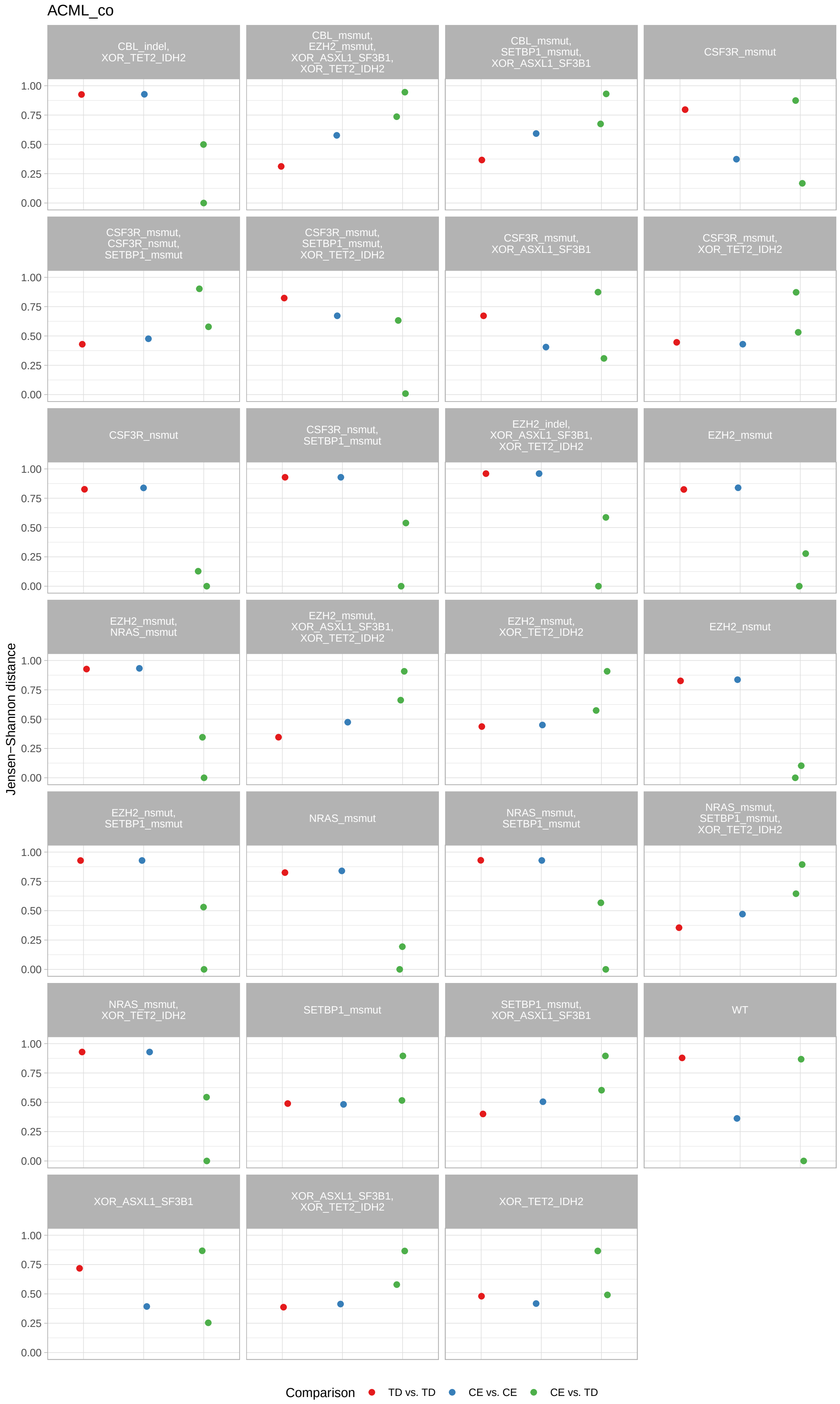
ACML_co
Jensen−Shannon distance
CBL_indel, XOR_TET2_IDH2
CBL_msmut, EZH2_msmut, XOR_ASXL1_SF3B1, XOR_TET2_IDH2
CBL_msmut, SETBP1_msmut, XOR_ASXL1_SF3B1
CSF3R_msmut
CSF3R_msmut, CSF3R_nsmut, SETBP1_msmut
CSF3R_msmut, SETBP1_msmut, XOR_TET2_IDH2
CSF3R_msmut, XOR_ASXL1_SF3B1
CSF3R_msmut, XOR_TET2_IDH2
CSF3R_nsmut
CSF3R_nsmut, SETBP1_msmut
EZH2_indel, XOR_ASXL1_SF3B1, XOR_TET2_IDH2
EZH2_msmut
EZH2_msmut, NRAS_msmut
EZH2_msmut, XOR_ASXL1_SF3B1, XOR_TET2_IDH2
EZH2_msmut, XOR_TET2_IDH2
EZH2_nsmut
EZH2_nsmut, SETBP1_msmut
NRAS_msmut
NRAS_msmut, SETBP1_msmut
NRAS_msmut, SETBP1_msmut, XOR_TET2_IDH2
NRAS_msmut, XOR_TET2_IDH2
SETBP1_msmut
SETBP1_msmut, XOR_ASXL1_SF3B1
WT
XOR_ASXL1_SF3B1
XOR_ASXL1_SF3B1, XOR_TET2_IDH2
XOR_TET2_IDH2
1.00
0.75
0.50
0.25
0.00
Comparison
TD vs. TD
CE vs. CE
CE vs. TD

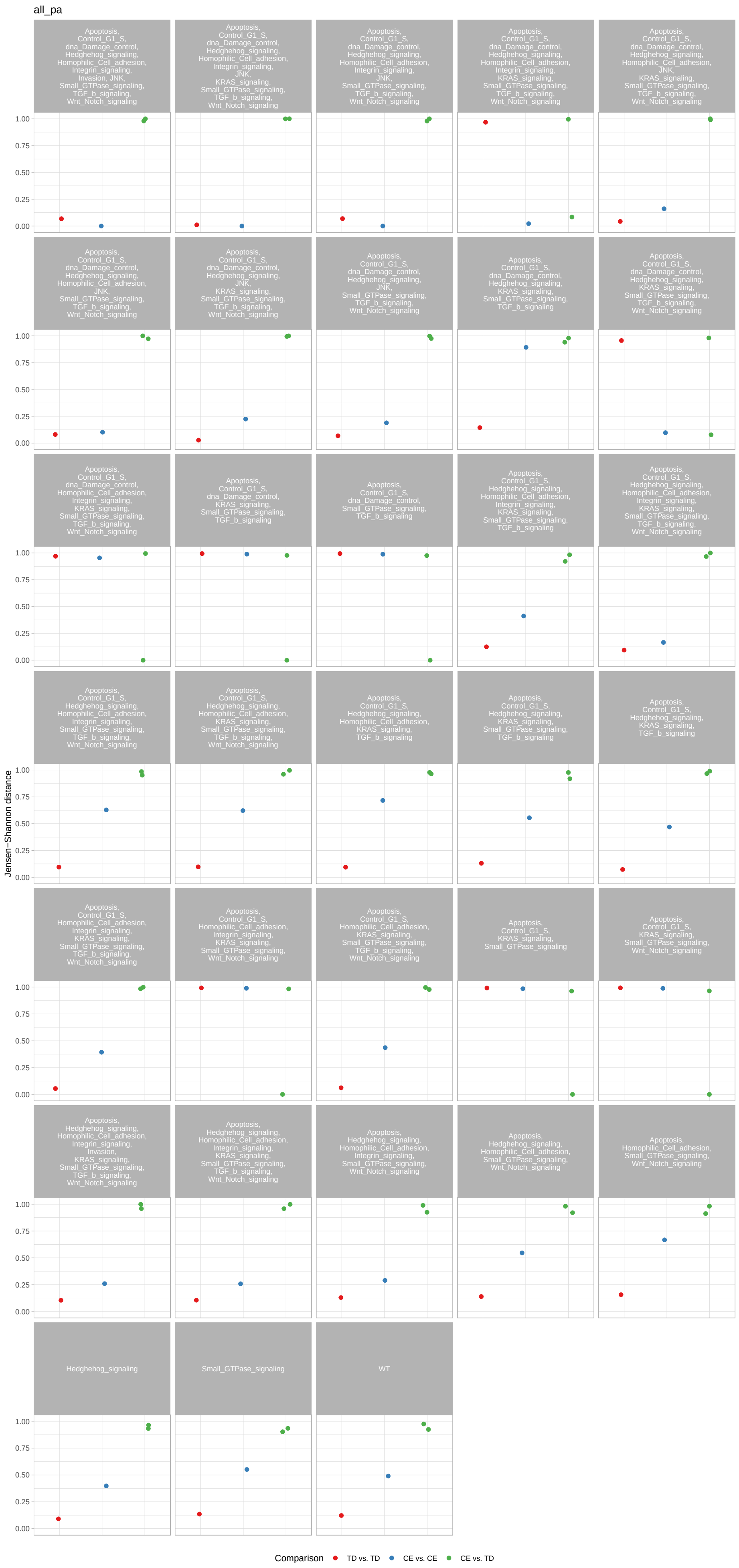
all_pa
Jensen−Shannon distance
Comparison
TD vs. TD
CE vs. CE
CE vs. TD
Hedghehog_signaling
Small_GTPase_signaling
WT

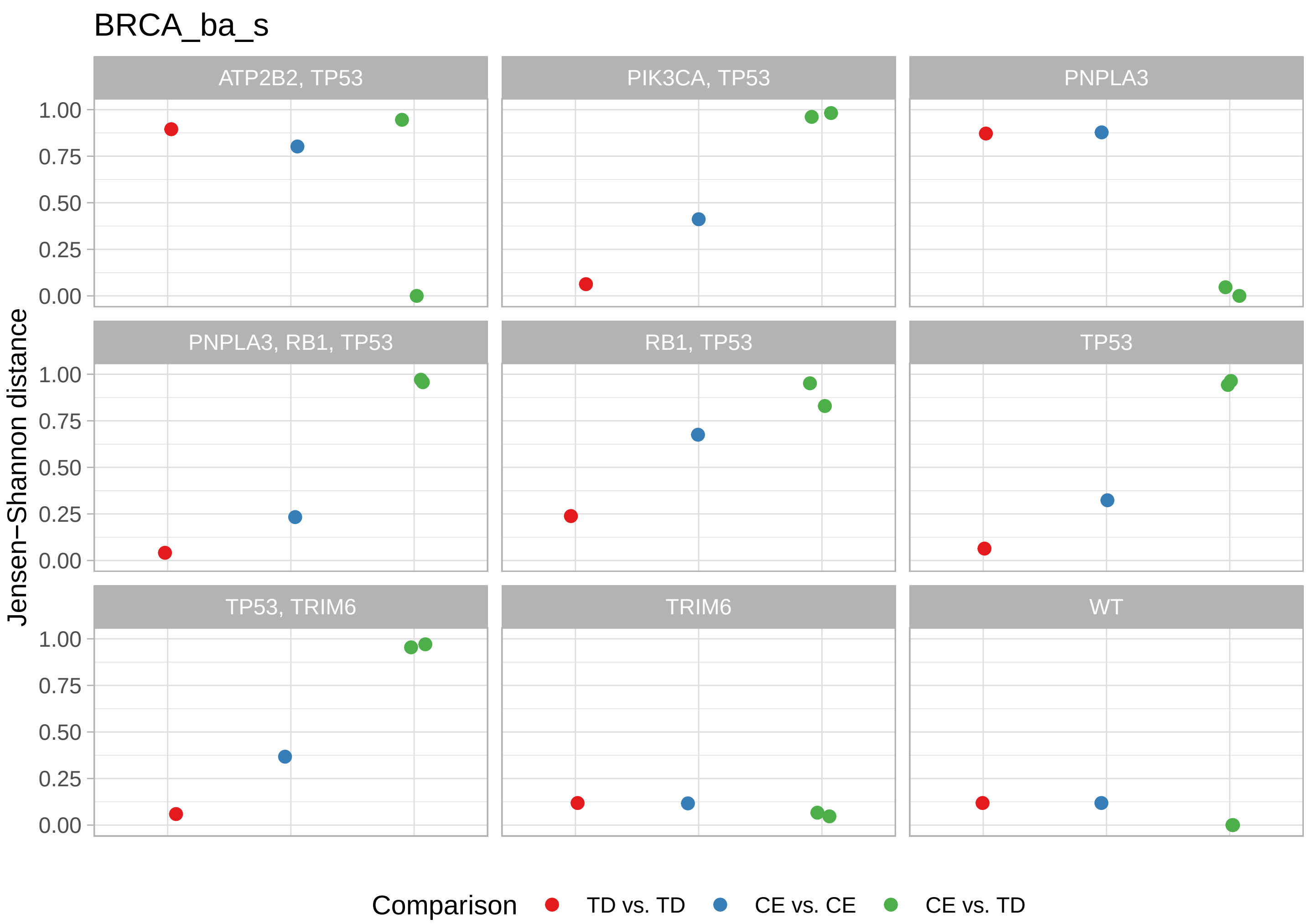

BRCA_ba_s
ATP2B2, TP53
PIK3CA, TP53
PNPLA3
PNPLA3, RB1, TP53
RB1, TP53
TP53
TP53, TRIM6
TRIM6
WT
Jensen−Shannon distance
1.00
0.75
0.50
0.25
0.00
Comparison
TD vs. TD
CE vs. CE
CE vs. TD

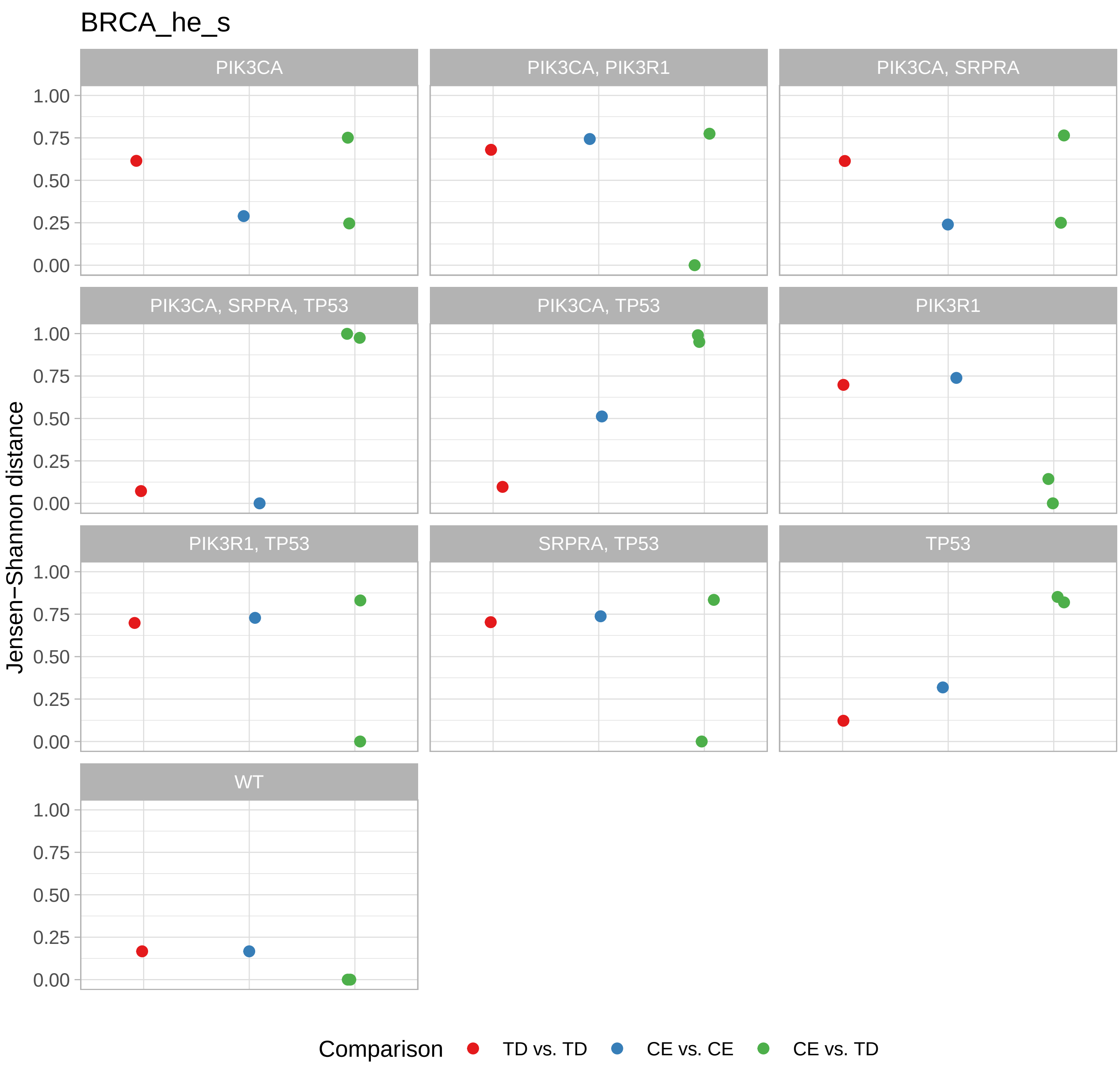

BRCA_he_s
PIK3CA
PIK3CA, PIK3R1
PIK3CA, SRPRA
PIK3CA, SRPRA, TP53
PIK3CA, TP53
PIK3R1
PIK3R1, TP53
SRPRA, TP53
TP53
WT
1.00
0.75
0.50
0.25
0.00
Jensen−Shannon distance
Comparison
TD vs. TD
CE vs. CE
CE vs. TD

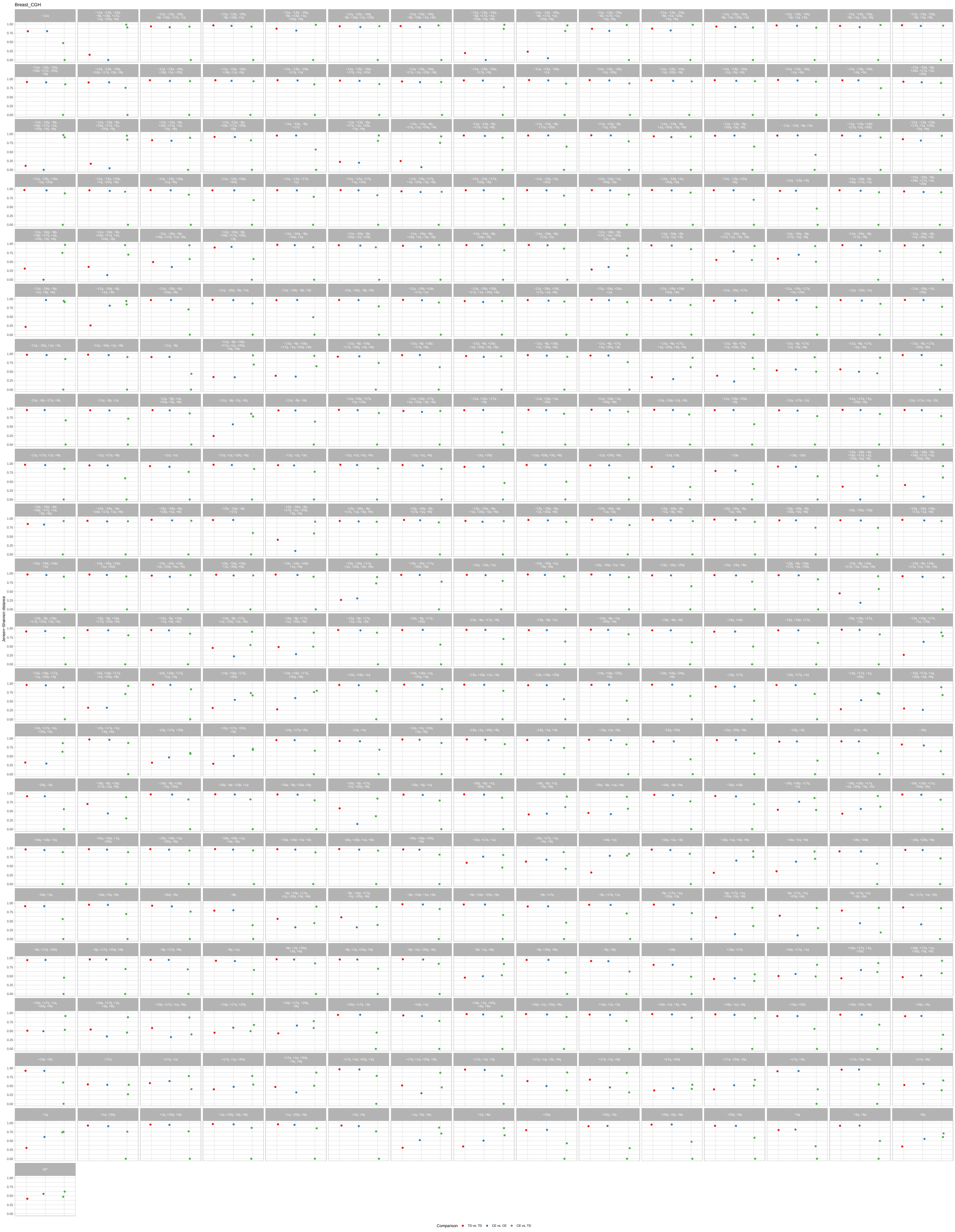
Breast_CGH
Jensen–Shannon distance
Comparison
TD vs. TD
CE vs. CE
CE vs. TD

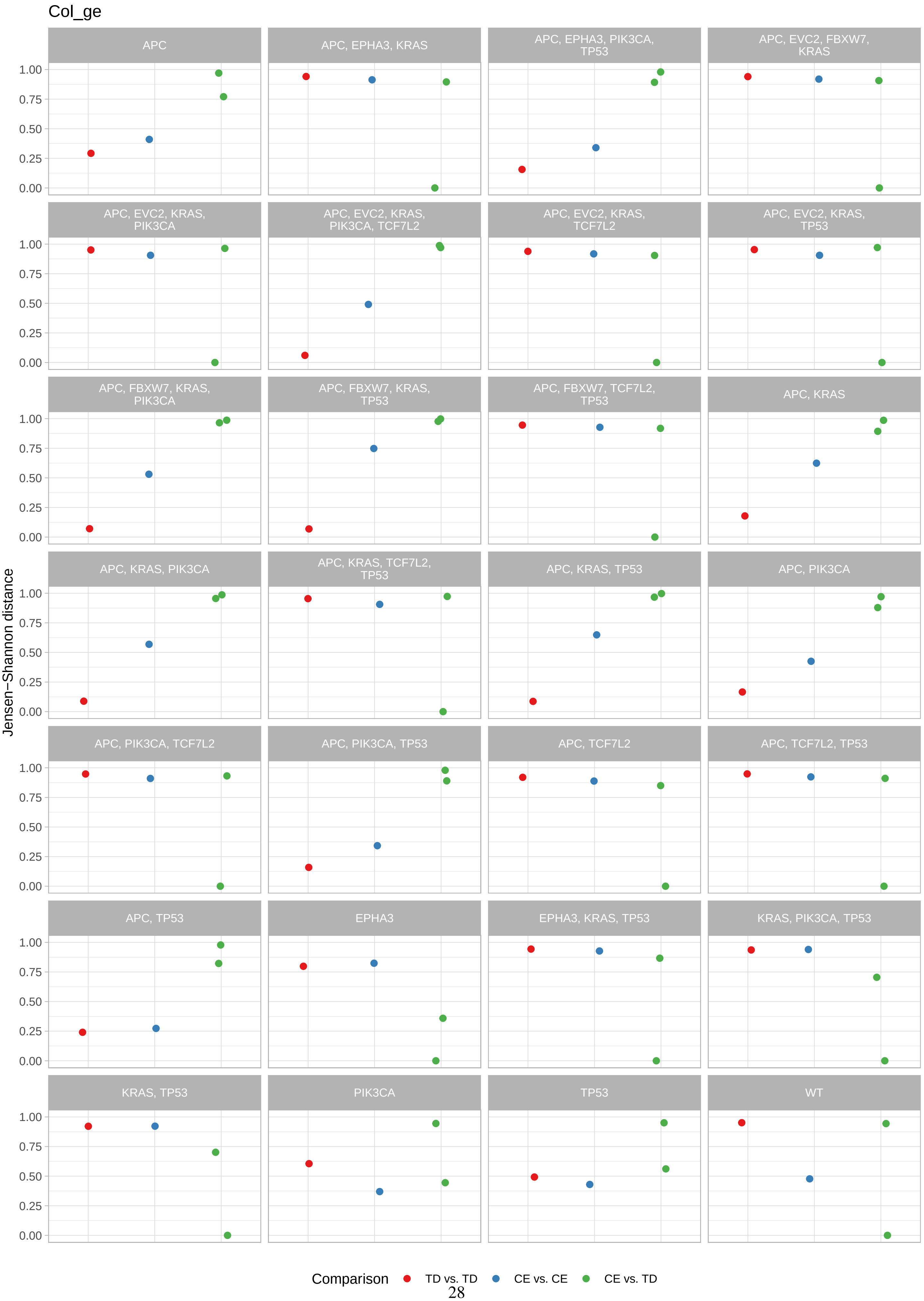

Col_ge
APC
APC, EPHA3, KRAS
APC, EPHA3, PIK3CA, TP53
APC, EVC2, FBXW7, KRAS
APC, EVC2, KRAS, PIK3CA
APC, EVC2, KRAS, PIK3CA, TCF7L2
APC, EVC2, KRAS, TCF7L2
APC, EVC2, KRAS, TP53
APC, FBXW7, KRAS, PIK3CA
APC, FBXW7, KRAS, TP53
APC, FBXW7, TCF7L2, TP53
APC, KRAS
APC, KRAS, PIK3CA
APC, KRAS, TCF7L2, TP53
APC, KRAS, TP53
APC, PIK3CA
APC, PIK3CA, TCF7L2
APC, PIK3CA, TP53
APC, TCF7L2
APC, TCF7L2, TP53
APC, TP53
EPHA3
EPHA3, KRAS, TP53
KRAS, PIK3CA, TP53
KRAS, TP53
PIK3CA
TP53
WT
Jensen−Shannon distance
0.00
0.25
0.50
0.75
1.00
Comparison
TD vs. TD
CE vs. CE
CE vs. TD

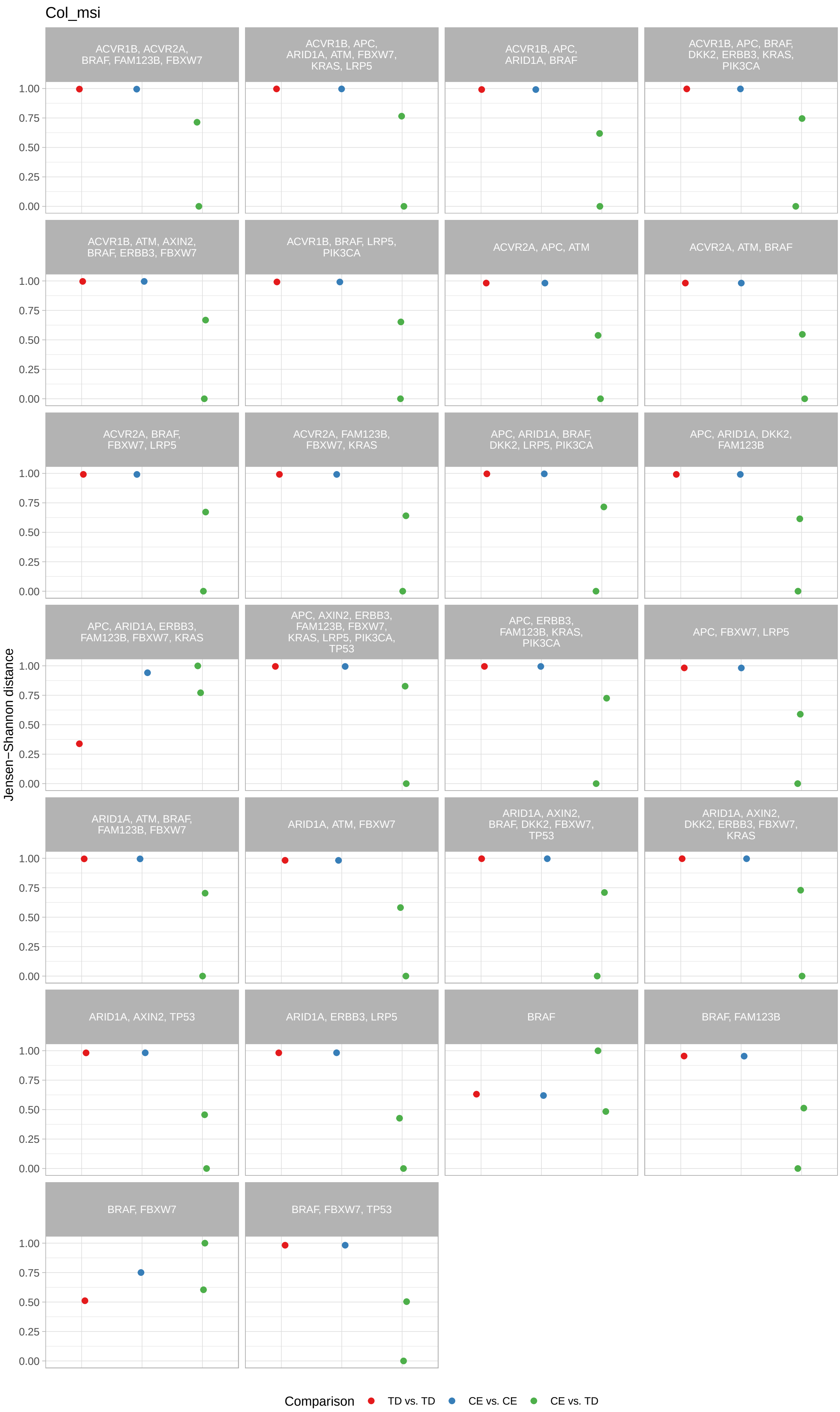
Col_msi
ACVR1B, ACVR2A, BRAF, FAM123B, FBXW7
ACVR1B, APC, ARID1A, ATM, FBXW7, KRAS, LRP5
ACVR1B, APC, ARID1A, BRAF
ACVR1B, APC, BRAF, DKK2, ERBB3, KRAS, PIK3CA
ACVR1B, ATM, AXIN2, BRAF, ERBB3, FBXW7
ACVR1B, BRAF, LRP5, PIK3CA
ACVR2A, APC, ATM
ACVR2A, ATM, BRAF
ACVR2A, BRAF, FBXW7, LRP5
ACVR2A, FAM123B, FBXW7, KRAS
APC, ARID1A, BRAF, DKK2, LRP5, PIK3CA
APC, ARID1A, DKK2, FAM123B
APC, ARID1A, ERBB3, FAM123B, FBXW7, KRAS
APC, AXIN2, ERBB3, FAM123B, FBXW7, KRAS, LRP5, PIK3CA, TP53
APC, ERBB3, FAM123B, KRAS, PIK3CA
APC, FBXW7, LRP5
ARID1A, ATM, BRAF, FAM123B, FBXW7
ARID1A, ATM, FBXW7
ARID1A, AXIN2, BRAF, DKK2, FBXW7, TP53
ARID1A, AXIN2, DKK2, ERBB3, FBXW7, KRAS
ARID1A, AXIN2, TP53
ARID1A, ERBB3, LRP5
BRAF
BRAF, FAM123B
BRAF, FBXW7
BRAF, FBXW7, TP53
Jensen−Shannon distance
1.00
0.75
0.50
0.25
0.00
Comparison
TD vs. TD
CE vs. CE
CE vs. TD

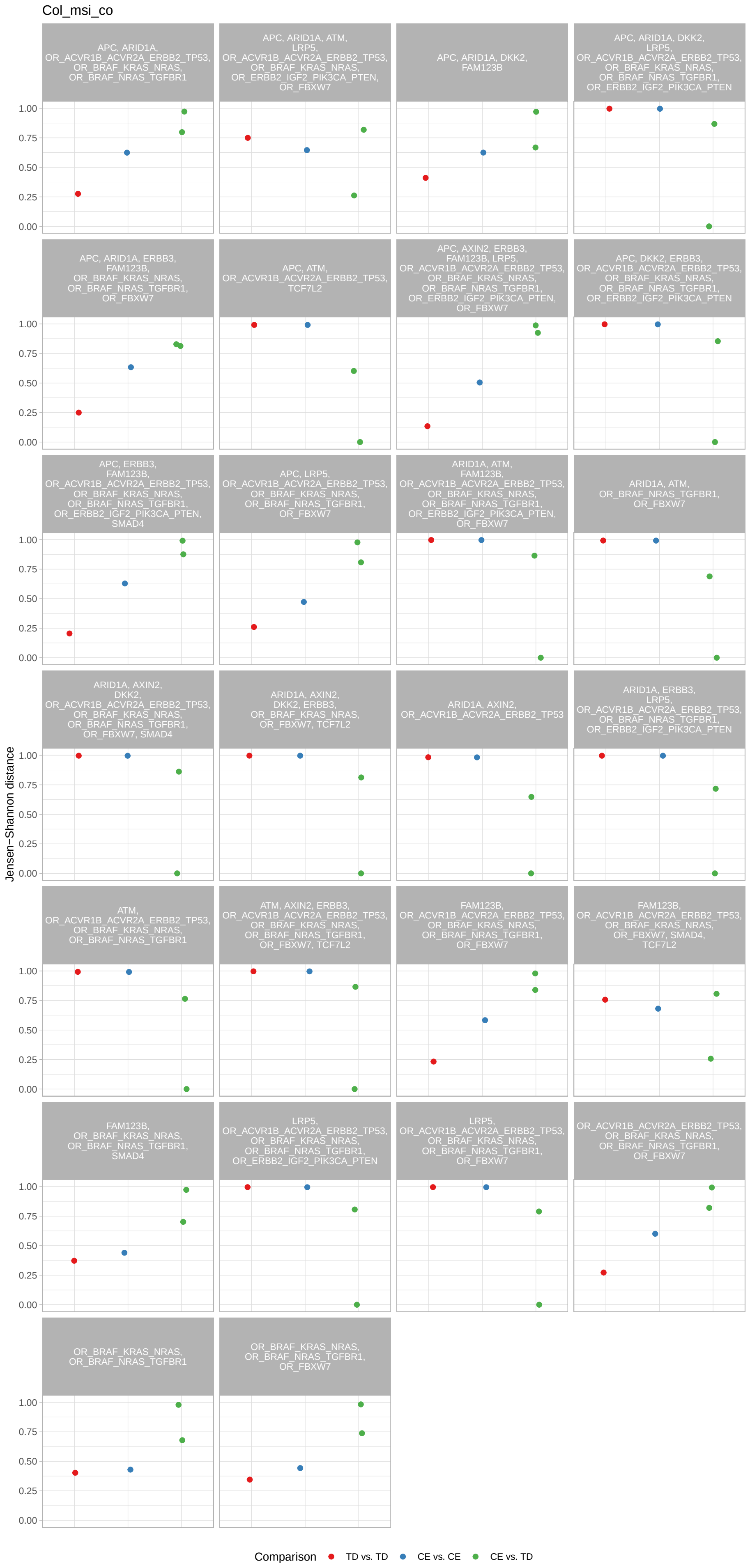
Col_msi_co
APC, ARID1A, OR_ACVR1B_ACVR2A_ERBB2_TP53, OR_BRAF_KRAS_NRAS, OR_BRAF_NRAS_TGFBR1
APC, ARID1A, ATM, LRP5, OR_ACVR1B_ACVR2A_ERBB2_TP53, OR_BRAF_KRAS_NRAS, OR_ERBB2_IGF2_PIK3CA_PTEN, OR_FBXW7
APC, ARID1A, DKK2, FAM123B
APC, ARID1A, DKK2, LRP5, OR_ACVR1B_ACVR2A_ERBB2_TP53, OR_BRAF_KRAS_NRAS, OR_BRAF_NRAS_TGFBR1, OR_ERBB2_IGF2_PIK3CA_PTEN
APC, ARID1A, ERBB3, FAM123B, OR_BRAF_KRAS_NRAS, OR_BRAF_NRAS_TGFBR1, OR_FBXW7
APC, ATM, OR_ACVR1B_ACVR2A_ERBB2_TP53, TCF7L2
APC, AXIN2, ERBB3, FAM123B, LRP5, OR_ACVR1B_ACVR2A_ERBB2_TP53, OR_BRAF_KRAS_NRAS, OR_BRAF_NRAS_TGFBR1, OR_ERBB2_IGF2_PIK3CA_PTEN, OR_FBXW7
APC, DKK2, ERBB3, OR_ACVR1B_ACVR2A_ERBB2_TP53, OR_BRAF_KRAS_NRAS, OR_BRAF_NRAS_TGFBR1, OR_ERBB2_IGF2_PIK3CA_PTEN
APC, ERBB3, FAM123B, OR_ACVR1B_ACVR2A_ERBB2_TP53, OR_BRAF_KRAS_NRAS, OR_BRAF_NRAS_TGFBR1, OR_ERBB2_IGF2_PIK3CA_PTEN, SMAD4
APC, LRP5, OR_ACVR1B_ACVR2A_ERBB2_TP53, OR_BRAF_KRAS_NRAS, OR_BRAF_NRAS_TGFBR1, OR_FBXW7
ARID1A, ATM, FAM123B, OR_ACVR1B_ACVR2A_ERBB2_TP53, OR_BRAF_KRAS_NRAS, OR_BRAF_NRAS_TGFBR1, OR_ERBB2_IGF2_PIK3CA_PTEN, OR_FBXW7
ARID1A, ATM, OR_BRAF_NRAS_TGFBR1, OR_FBXW7
ARID1A, AXIN2, DKK2, OR_ACVR1B_ACVR2A_ERBB2_TP53, OR_BRAF_KRAS_NRAS, OR_BRAF_NRAS_TGFBR1, OR_FBXW7, SMAD4
ARID1A, AXIN2, DKK2, ERBB3, OR_BRAF_KRAS_NRAS, OR_FBXW7, TCF7L2
ARID1A, AXIN2, OR_ACVR1B_ACVR2A_ERBB2_TP53
ARID1A, ERBB3, LRP5, OR_ACVR1B_ACVR2A_ERBB2_TP53, OR_BRAF_NRAS_TGFBR1, OR_ERBB2_IGF2_PIK3CA_PTEN
ATM, OR_ACVR1B_ACVR2A_ERBB2_TP53, OR_BRAF_KRAS_NRAS, OR_BRAF_NRAS_TGFBR1
ATM, AXIN2, ERBB3, OR_ACVR1B_ACVR2A_ERBB2_TP53, OR_BRAF_KRAS_NRAS, OR_BRAF_NRAS_TGFBR1, OR_FBXW7, TCF7L2
FAM123B, OR_ACVR1B_ACVR2A_ERBB2_TP53, OR_BRAF_KRAS_NRAS, OR_BRAF_NRAS_TGFBR1, OR_FBXW7
FAM123B, OR_ACVR1B_ACVR2A_ERBB2_TP53, OR_BRAF_KRAS_NRAS, OR_FBXW7, SMAD4, TCF7L2
FAM123B, OR_BRAF_KRAS_NRAS, OR_BRAF_NRAS_TGFBR1, SMAD4
LRP5, OR_ACVR1B_ACVR2A_ERBB2_TP53, OR_BRAF_KRAS_NRAS, OR_BRAF_NRAS_TGFBR1, OR_ERBB2_IGF2_PIK3CA_PTEN
LRP5, OR_ACVR1B_ACVR2A_ERBB2_TP53, OR_BRAF_KRAS_NRAS, OR_BRAF_NRAS_TGFBR1, OR_FBXW7
OR_ACVR1B_ACVR2A_ERBB2_TP53, OR_BRAF_KRAS_NRAS, OR_BRAF_NRAS_TGFBR1, OR_FBXW7
OR_BRAF_KRAS_NRAS, OR_BRAF_NRAS_TGFBR1
OR_BRAF_KRAS_NRAS, OR_BRAF_NRAS_TGFBR1, OR_FBXW7
Jensen−Shannon distance
1.00
0.75
0.50
0.25
0.00
Comparison
TD vs. TD
CE vs. CE
CE vs. TD

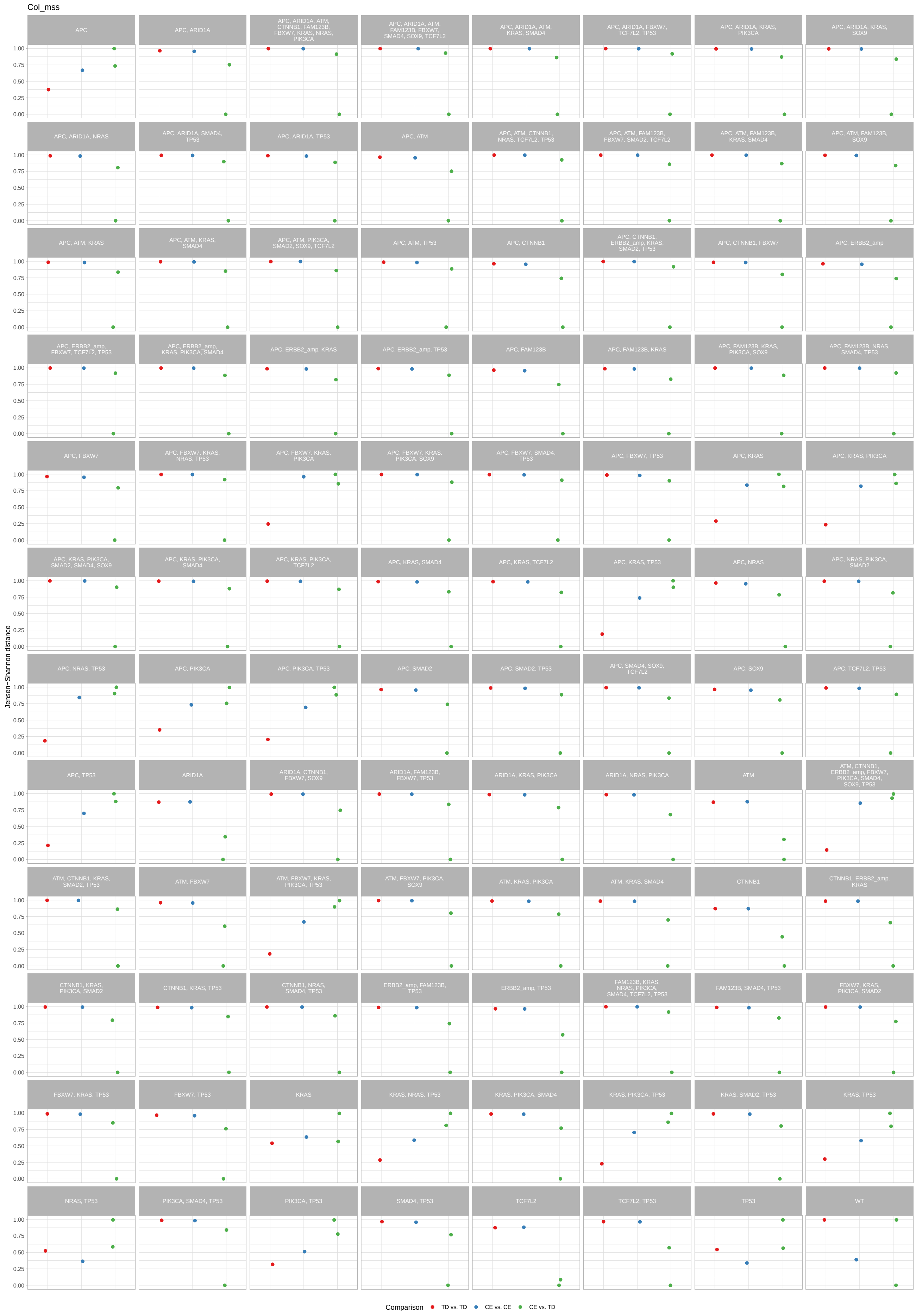

Col_mss
APC
APC, ARID1A
APC, ARID1A, ATM, CTNNB1, FAM123B, FBXW7, KRAS, NRAS, PIK3CA
APC, ARID1A, ATM, FAM123B, FBXW7, SMAD4, SOX9, TCF7L2
APC, ARID1A, ATM, KRAS, SMAD4
APC, ARID1A, FBXW7, TCF7L2, TP53
APC, ARID1A, KRAS, PIK3CA
APC, ARID1A, KRAS, SOX9
APC, ARID1A, NRAS
APC, ARID1A, SMAD4, TP53
APC, ARID1A, TP53
APC, ATM
APC, ATM, CTNNB1, NRAS, TCF7L2, TP53
APC, ATM, FAM123B, FBXW7, SMAD2, TCF7L2
APC, ATM, FAM123B, KRAS, SMAD4
APC, ATM, FAM123B, SOX9
APC, ATM, KRAS
APC, ATM, KRAS, SMAD4
APC, ATM, PIK3CA, SMAD2, SOX9, TCF7L2
APC, ATM, TP53
APC, CTNNB1
APC, CTNNB1, ERBB2_amp, KRAS, SMAD2, TP53
APC, CTNNB1, FBXW7
APC, ERBB2_amp
APC, ERBB2_amp, FBXW7, TCF7L2, TP53
APC, ERBB2_amp, KRAS, PIK3CA, SMAD4
APC, ERBB2_amp, KRAS
APC, ERBB2_amp, TP53
APC, FAM123B
APC, FAM123B, KRAS
APC, FAM123B, KRAS, PIK3CA, SOX9
APC, FAM123B, NRAS, SMAD4, TP53
APC, FBXW7
APC, FBXW7, KRAS, NRAS, TP53
APC, FBXW7, KRAS, PIK3CA
APC, FBXW7, KRAS, PIK3CA, SOX9
APC, FBXW7, SMAD4, TP53
APC, FBXW7, TP53
APC, KRAS
APC, KRAS, PIK3CA
APC, KRAS, PIK3CA, SMAD2, SMAD4, SOX9
APC, KRAS, PIK3CA, SMAD4
APC, KRAS, PIK3CA, TCF7L2
APC, KRAS, SMAD4
APC, KRAS, TCF7L2
APC, KRAS, TP53
APC, NRAS
APC, NRAS, PIK3CA, SMAD2
APC, NRAS, TP53
APC, PIK3CA
APC, PIK3CA, TP53
APC, SMAD2
APC, SMAD2, TP53
APC, SMAD4, SOX9, TCF7L2
APC, SOX9
APC, TCF7L2, TP53
APC, TP53
ARID1A
ARID1A, CTNNB1, FBXW7, SOX9
ARID1A, FAM123B, FBXW7, TP53
ARID1A, KRAS, PIK3CA
ARID1A, NRAS, PIK3CA
ATM
ATM, CTNNB1, ERBB2_amp, FBXW7, PIK3CA, SMAD4, SOX9, TP53
ATM, CTNNB1, KRAS, SMAD2, TP53
ATM, FBXW7
ATM, FBXW7, KRAS, PIK3CA, TP53
ATM, FBXW7, PIK3CA, SOX9
ATM, KRAS, PIK3CA
ATM, KRAS, SMAD4
CTNNB1
CTNNB1, ERBB2_amp, KRAS
CTNNB1, KRAS, PIK3CA, SMAD2
CTNNB1, KRAS, TP53
CTNNB1, NRAS, SMAD4, TP53
ERBB2_amp, FAM123B, TP53
ERBB2_amp, TP53
FAM123B, KRAS, NRAS, PIK3CA, SMAD4, TCF7L2, TP53
FAM123B, SMAD4, TP53
FBXW7, KRAS, PIK3CA, SMAD2
FBXW7, KRAS, TP53
FBXW7, TP53
KRAS
KRAS, NRAS, TP53
KRAS, PIK3CA, SMAD4
KRAS, PIK3CA, TP53
KRAS, SMAD2, TP53
KRAS, TP53
NRAS, TP53
PIK3CA, SMAD4, TP53
PIK3CA, TP53
SMAD4, TP53
TCF7L2
TCF7L2, TP53
TP53
WT
Jensen−Shannon distance
1.00
0.75
0.50
0.25
0.00
Comparison
TD vs. TD
CE vs. CE
CE vs. TD

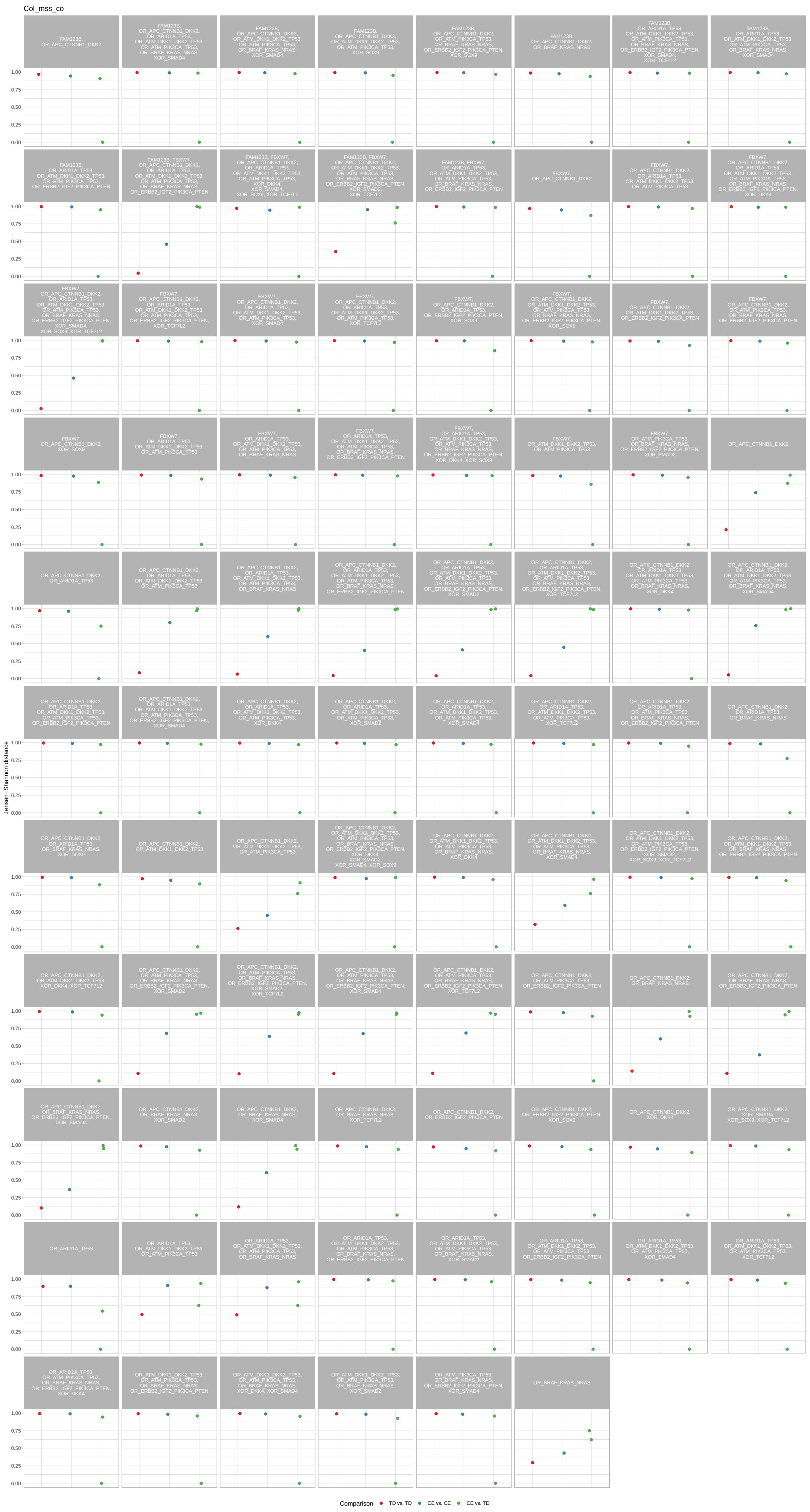
Col_mss_co
Jensen-Shannon distance
Comparison
TD vs. TD
CE vs. CE
CE vs. TD

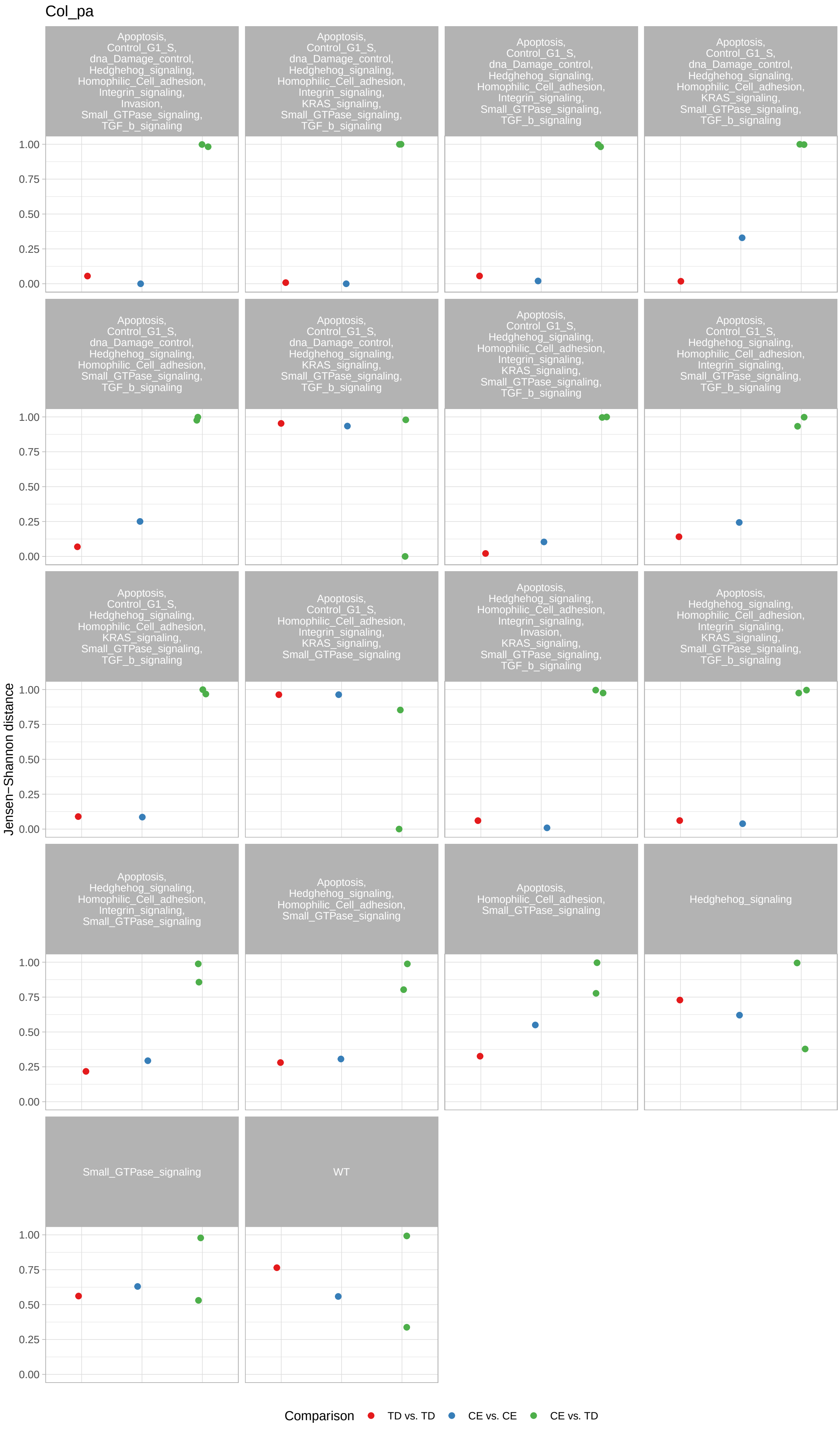
Col_pa
Apoptosis, Control_G1_S, dna_Damage_control, Hedghehog_signaling, Homophilic_Cell_adhesion, Integrin_signaling, Invasion, Small_GTPase_signaling, TGF_b_signaling
Apoptosis, Control_G1_S, dna_Damage_control, Hedghehog_signaling, Homophilic_Cell_adhesion, Integrin_signaling, KRAS_signaling, Small_GTPase_signaling, TGF_b_signaling
Apoptosis, Control_G1_S, dna_Damage_control, Hedghehog_signaling, Homophilic_Cell_adhesion, Integrin_signaling, Small_GTPase_signaling, TGF_b_signaling
Apoptosis, Control_G1_S, dna_Damage_control, Hedghehog_signaling, Homophilic_Cell_adhesion, KRAS_signaling, Small_GTPase_signaling, TGF_b_signaling
Apoptosis, Control_G1_S, dna_Damage_control, Hedghehog_signaling, Homophilic_Cell_adhesion, Small_GTPase_signaling, TGF_b_signaling
Apoptosis, Control_G1_S, dna_Damage_control, Hedghehog_signaling, KRAS_signaling, Small_GTPase_signaling, TGF_b_signaling
Apoptosis, Control_G1_S, Hedghehog_signaling, Homophilic_Cell_adhesion, Integrin_signaling, KRAS_signaling, Small_GTPase_signaling, TGF_b_signaling
Apoptosis, Control_G1_S, Hedghehog_signaling, Homophilic_Cell_adhesion, Integrin_signaling, Small_GTPase_signaling, TGF_b_signaling
Apoptosis, Control_G1_S, Hedghehog_signaling, Homophilic_Cell_adhesion, KRAS_signaling, Small_GTPase_signaling, TGF_b_signaling
Apoptosis, Control_G1_S, Homophilic_Cell_adhesion, Integrin_signaling, KRAS_signaling, Small_GTPase_signaling
Apoptosis, Hedghehog_signaling, Homophilic_Cell_adhesion, Integrin_signaling, Invasion, KRAS_signaling, Small_GTPase_signaling, TGF_b_signaling
Apoptosis, Hedghehog_signaling, Homophilic_Cell_adhesion, Integrin_signaling, KRAS_signaling, Small_GTPase_signaling, TGF_b_signaling
Apoptosis, Hedghehog_signaling, Homophilic_Cell_adhesion, Integrin_signaling, Small_GTPase_signaling
Apoptosis, Hedghehog_signaling, Homophilic_Cell_adhesion, Small_GTPase_signaling
Apoptosis, Homophilic_Cell_adhesion, Small_GTPase_signaling
Hedghehog_signaling
Small_GTPase_signaling
WT
Jensen−Shannon distance
0.00
0.25
0.50
0.75
1.00
Comparison
TD vs. TD
CE vs. CE
CE vs. TD

Colon_CGH

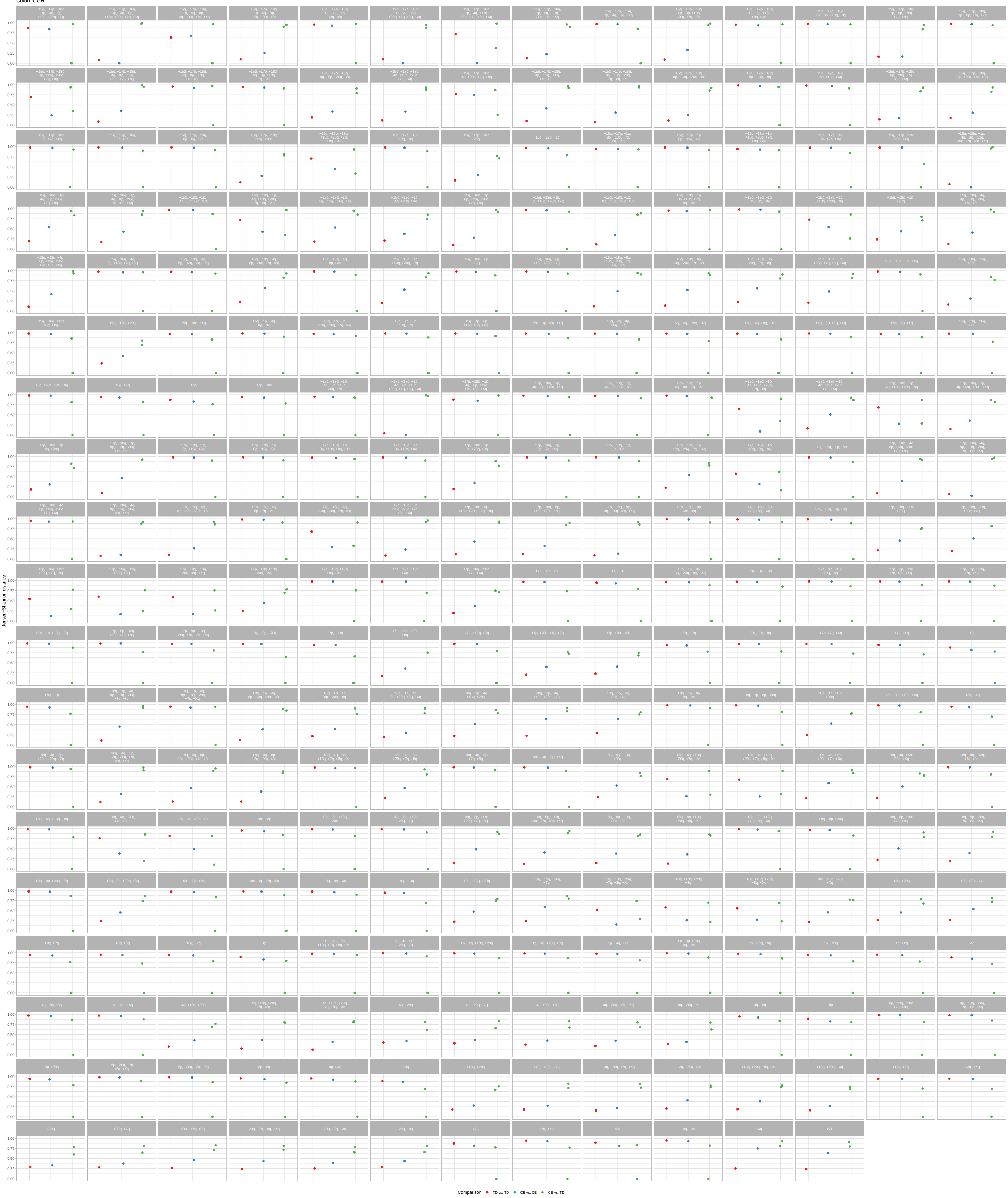

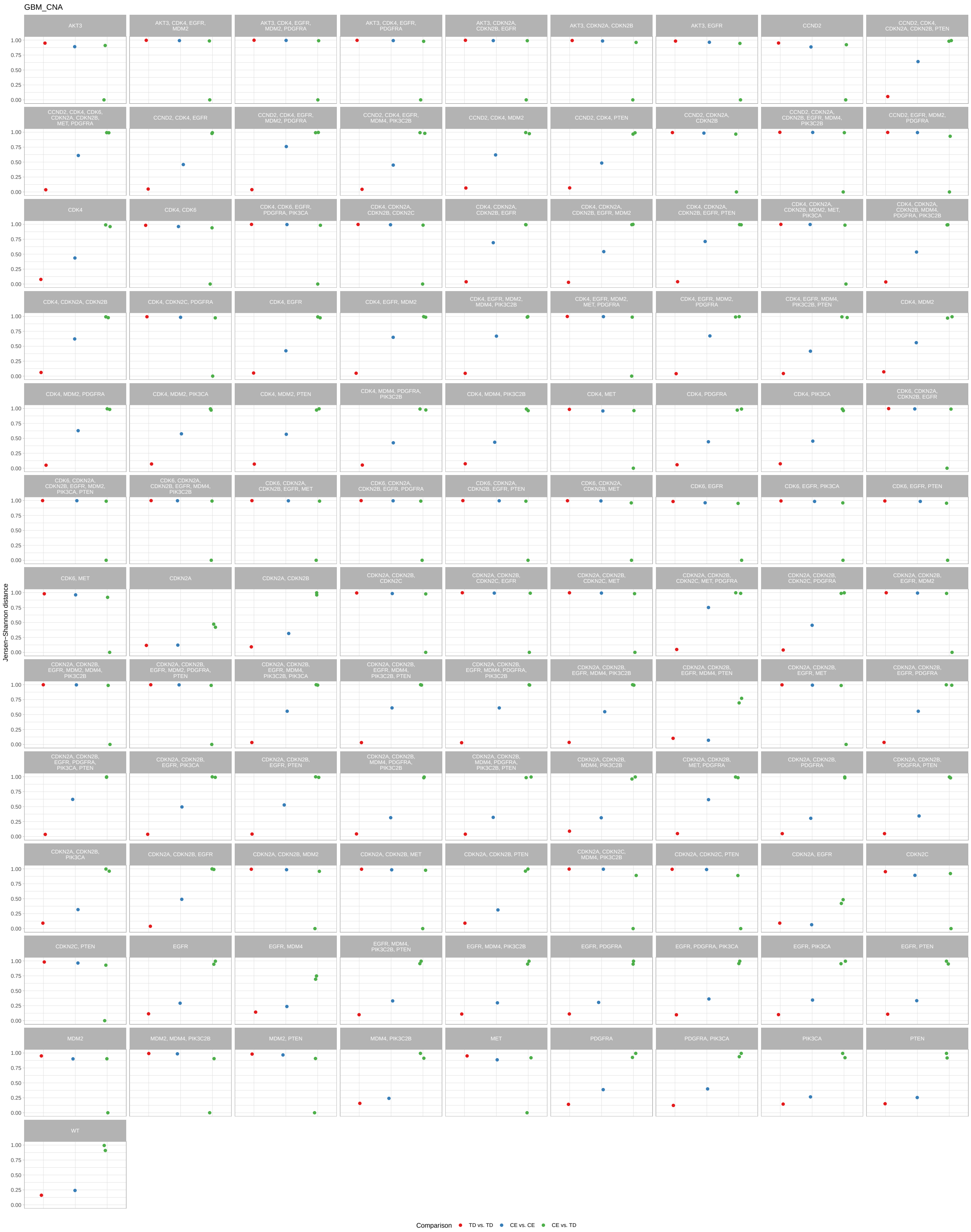
GBM_CNA
AKT3
AKT3, CDK4, EGFR, MDM2
AKT3, CDK4, EGFR, MDM2, PDGFRA
AKT3, CDK4, EGFR, PDGFRA
AKT3, CDKN2A, CDKN2B, EGFR
AKT3, CDKN2A, CDKN2B
AKT3, EGFR
CCND2
CCND2, CDK4, CDKN2A, CDKN2B, PTEN
CCND2, CDK4, CDK6, CDKN2A, CDKN2B, MET, PDGFRA
CCND2, CDK4, EGFR
CCND2, CDK4, EGFR, MDM2, PDGFRA
CCND2, CDK4, EGFR, MDM4, PIK3C2B
CCND2, CDK4, MDM2
CCND2, CDK4, PTEN
CCND2, CDKN2A, CDKN2B
CCND2, CDKN2A, CDKN2B, EGFR, MDM4, PIK3C2B
CCND2, EGFR, MDM2, PDGFRA
CDK4
CDK4, CDK6
CDK4, CDK6, EGFR, PDGFRA, PIK3CA
CDK4, CDKN2A, CDKN2B, CDKN2C
CDK4, CDKN2A, CDKN2B, EGFR
CDK4, CDKN2A, CDKN2B, EGFR, MDM2
CDK4, CDKN2A, CDKN2B, EGFR, PTEN
CDK4, CDKN2A, CDKN2B, MDM2, MET, PIK3CA
CDK4, CDKN2A, CDKN2B, MDM4, PDGFRA, PIK3C2B
CDK4, CDKN2A, CDKN2B
CDK4, CDKN2C, PDGFRA
CDK4, EGFR
CDK4, EGFR, MDM2
CDK4, EGFR, MDM2, MDM4, PIK3C2B
CDK4, EGFR, MDM2, MET, PDGFRA
CDK4, EGFR, MDM2, PDGFRA
CDK4, EGFR, MDM4, PIK3C2B, PTEN
CDK4, MDM2
CDK4, MDM2, PDGFRA
CDK4, MDM2, PIK3CA
CDK4, MDM2, PTEN
CDK4, MDM4, PDGFRA, PIK3C2B
CDK4, MDM4, PIK3C2B
CDK4, MET
CDK4, PDGFRA
CDK4, PIK3CA
CDK6, CDKN2A, CDKN2B, EGFR
CDK6, CDKN2A, CDKN2B, EGFR, MDM2, PIK3CA, PTEN
CDK6, CDKN2A, CDKN2B, EGFR, MDM4, PIK3C2B
CDK6, CDKN2A, CDKN2B, EGFR, MET
CDK6, CDKN2A, CDKN2B, EGFR, PDGFRA
CDK6, CDKN2A, CDKN2B, EGFR, PTEN
CDK6, CDKN2A, CDKN2B, MET
CDK6, EGFR
CDK6, EGFR, PIK3CA
CDK6, EGFR, PTEN
CDK6, MET
CDKN2A
CDKN2A, CDKN2B
CDKN2A, CDKN2B, CDKN2C
CDKN2A, CDKN2B, CDKN2C, EGFR
CDKN2A, CDKN2B, CDKN2C, MET
CDKN2A, CDKN2B, CDKN2C, MET, PDGFRA
CDKN2A, CDKN2B, CDKN2C, PDGFRA
CDKN2A, CDKN2B, EGFR, MDM2
CDKN2A, CDKN2B, EGFR, MDM2, MDM4, PIK3C2B
CDKN2A, CDKN2B, EGFR, MDM2, PDGFRA, PTEN
CDKN2A, CDKN2B, EGFR, MDM4, PIK3C2B, PIK3CA
CDKN2A, CDKN2B, EGFR, MDM4, PIK3C2B, PTEN
CDKN2A, CDKN2B, EGFR, MDM4, PDGFRA, PIK3C2B
CDKN2A, CDKN2B, EGFR, MDM4, PIK3C2B
CDKN2A, CDKN2B, EGFR, MDM4, PTEN
CDKN2A, CDKN2B, EGFR, MET
CDKN2A, CDKN2B, EGFR, PDGFRA
CDKN2A, CDKN2B, EGFR, PDGFRA, PIK3CA, PTEN
CDKN2A, CDKN2B, EGFR, PIK3CA
CDKN2A, CDKN2B, EGFR, PTEN
CDKN2A, CDKN2B, MDM4, PDGFRA, PIK3C2B
CDKN2A, CDKN2B, MDM4, PDGFRA, PIK3C2B, PTEN
CDKN2A, CDKN2B, MDM4, PIK3C2B
CDKN2A, CDKN2B, MET, PDGFRA
CDKN2A, CDKN2B, PDGFRA
CDKN2A, CDKN2B, PDGFRA, PTEN
CDKN2A, CDKN2B, PIK3CA
CDKN2A, CDKN2B, EGFR
CDKN2A, CDKN2B, MDM2
CDKN2A, CDKN2B, MET
CDKN2A, CDKN2B, PTEN
CDKN2A, CDKN2C, MDM4, PIK3C2B
CDKN2A, CDKN2C, PTEN
CDKN2A, EGFR
CDKN2C
CDKN2C, PTEN
EGFR
EGFR, MDM4
EGFR, MDM4, PIK3C2B, PTEN
EGFR, MDM4, PIK3C2B
EGFR, PDGFRA
EGFR, PDGFRA, PIK3CA
EGFR, PIK3CA
EGFR, PTEN
MDM2
MDM2, MDM4, PIK3C2B
MDM2, PTEN
MDM4, PIK3C2B
MET
PDGFRA
PDGFRA, PIK3CA
PIK3CA
PTEN
WT
1.00
0.75
0.50
0.25
0.00
Jensen−Shannon distance
Comparison
TD vs. TD
CE vs. CE
CE vs. TD

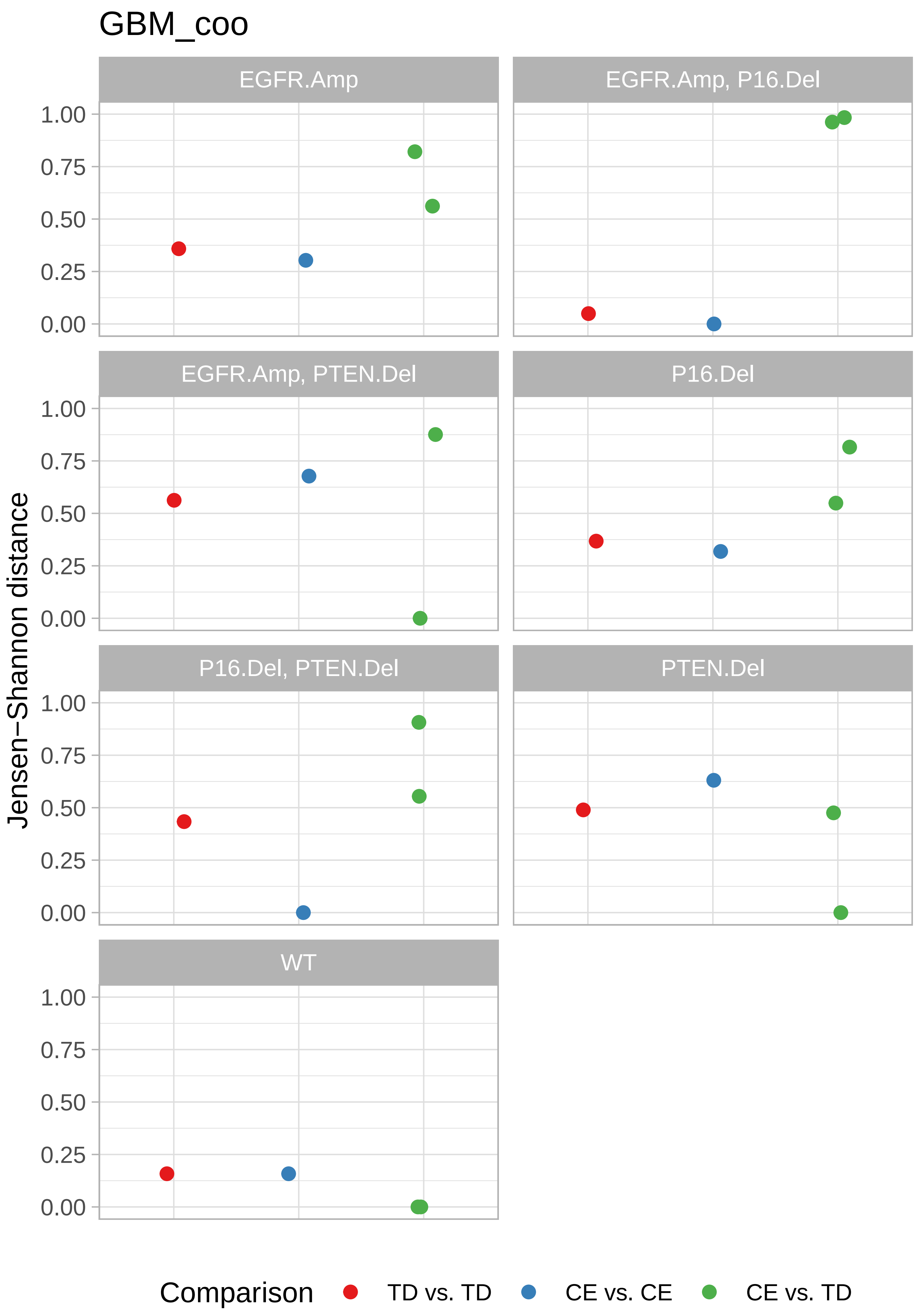
GBM_coo
EGFR.Amp
EGFR.Amp, P16.Del
EGFR.Amp, PTEN.Del
P16.Del
P16.Del, PTEN.Del
PTEN.Del
WT
Jensen−Shannon distance
1.00
0.75
0.50
0.25
0.00
Comparison
TD vs. TD
CE vs. CE
CE vs. TD

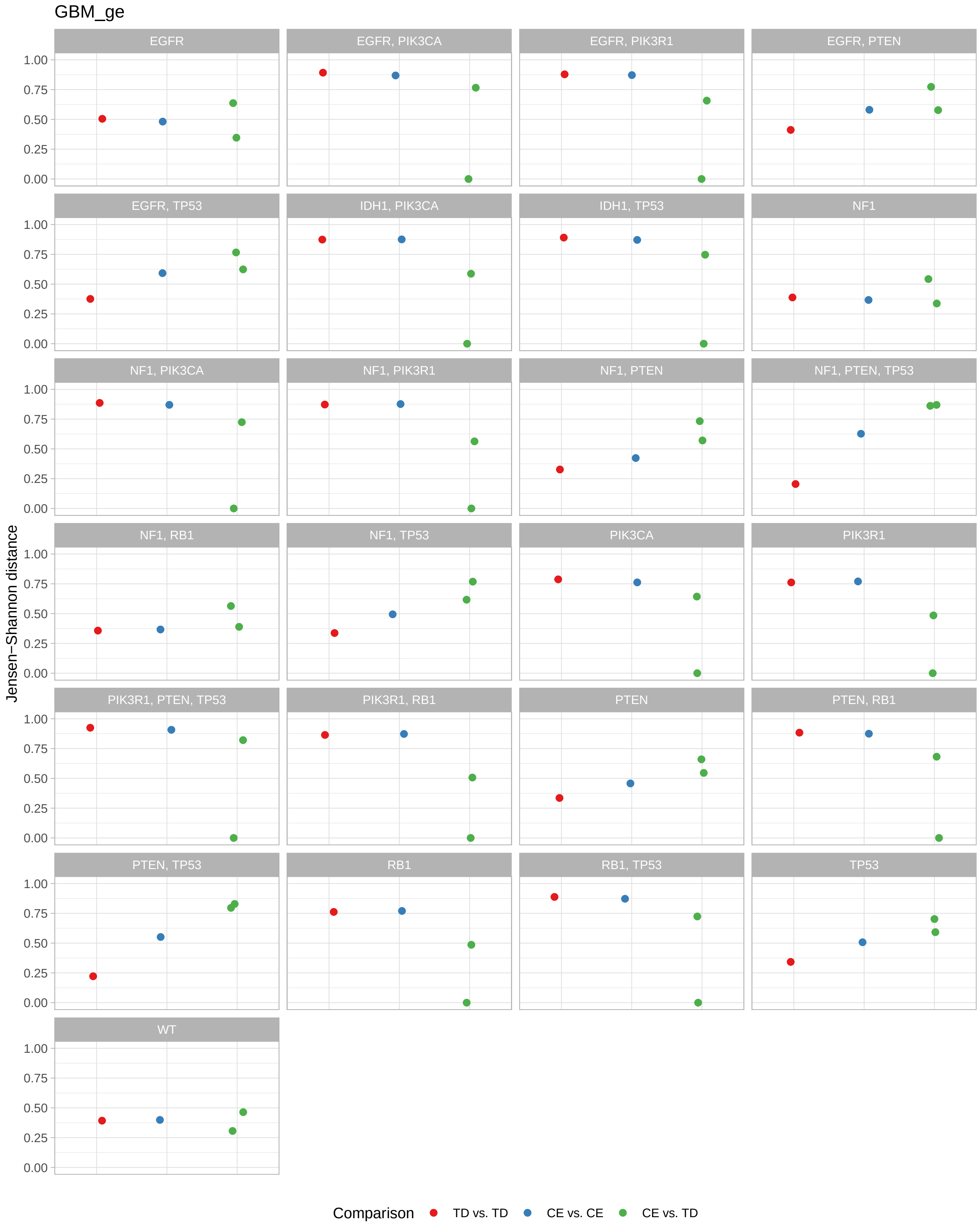
GBM_ge
EGFR
EGFR, PIK3CA
EGFR, PIK3R1
EGFR, PTEN
EGFR, TP53
IDH1, PIK3CA
IDH1, TP53
NF1
NF1, PIK3CA
NF1, PIK3R1
NF1, PTEN
NF1, PTEN, TP53
NF1, RB1
NF1, TP53
PIK3CA
PIK3R1
PIK3R1, PTEN, TP53
PIK3R1, RB1
PTEN
PTEN, RB1
PTEN, TP53
RB1
RB1, TP53
TP53
WT
1.00
0.75
0.50
0.25
0.00
Jensen−Shannon distance
Comparison
TD vs. TD
CE vs. CE
CE vs. TD

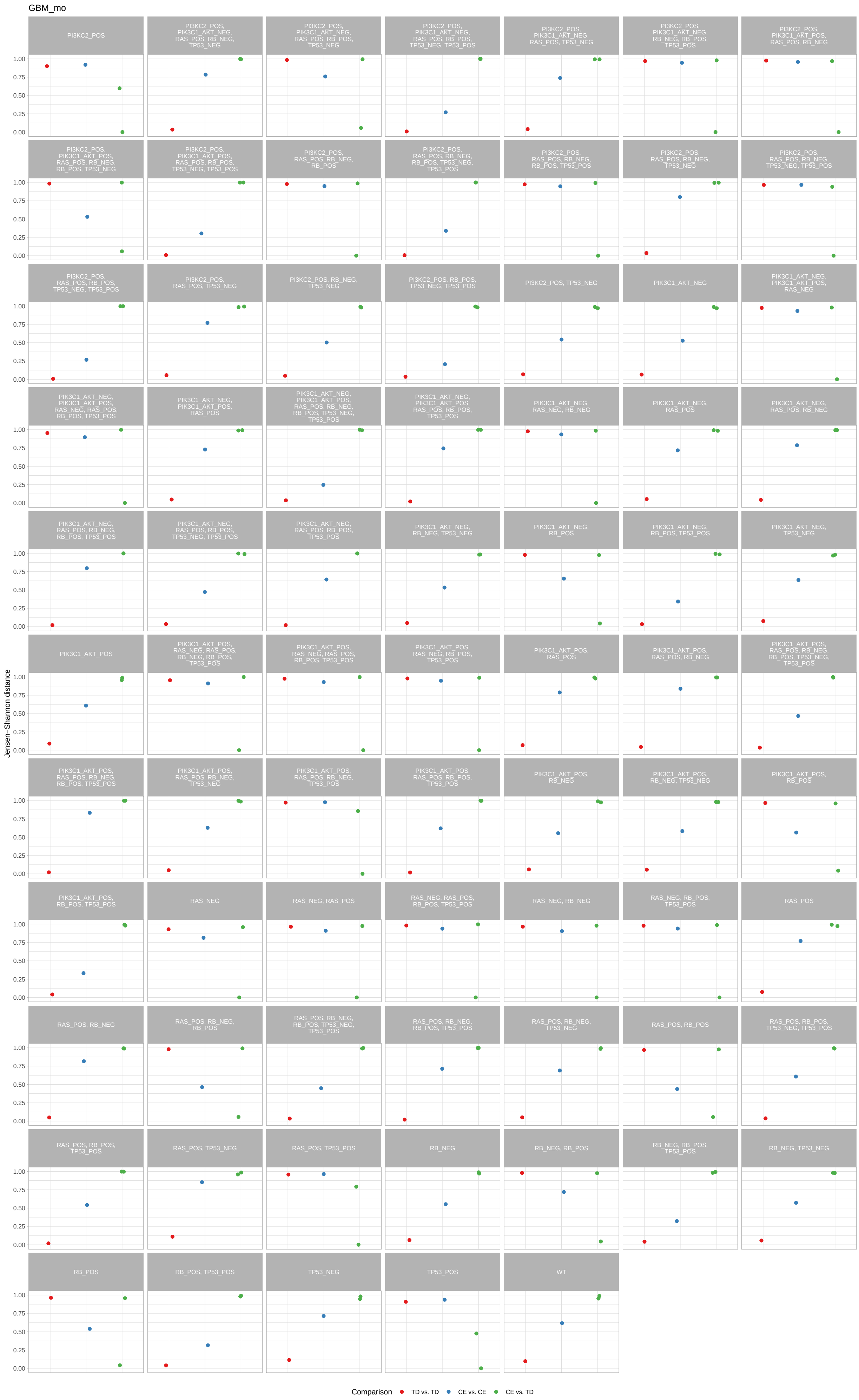
GBM_mo
PI3KC2_POS
PI3KC2_POS, PIK3C1_AKT_NEG, RAS_POS, RB_NEG, TP53_NEG
PI3KC2_POS, PIK3C1_AKT_NEG, RAS_POS, RB_POS, TP53_NEG
PI3KC2_POS, PIK3C1_AKT_NEG, RAS_POS, RB_POS, TP53_NEG, TP53_POS
PI3KC2_POS, PIK3C1_AKT_NEG, RAS_POS, TP53_NEG
PI3KC2_POS, PIK3C1_AKT_NEG, RB_NEG, RB_POS, TP53_POS
PI3KC2_POS, PIK3C1_AKT_POS, RAS_POS, RB_NEG
PI3KC2_POS, PIK3C1_AKT_POS, RAS_POS, RB_NEG, RB_POS, TP53_NEG
PI3KC2_POS, PIK3C1_AKT_POS, RAS_POS, RB_POS, TP53_NEG, TP53_POS
PI3KC2_POS, RAS_POS, RB_NEG, RB_POS
PI3KC2_POS, RAS_POS, RB_NEG, RB_POS, TP53_NEG, TP53_POS
PI3KC2_POS, RAS_POS, RB_NEG, RB_POS, TP53_POS
PI3KC2_POS, RAS_POS, RB_NEG, TP53_NEG
PI3KC2_POS, RAS_POS, RB_NEG, TP53_NEG, TP53_POS
PI3KC2_POS, RAS_POS, RB_POS, TP53_NEG, TP53_POS
PI3KC2_POS, RAS_POS, TP53_NEG
PI3KC2_POS, RB_NEG, TP53_NEG
PI3KC2_POS, RB_POS, TP53_NEG, TP53_POS
PI3KC2_POS, TP53_NEG
PIK3C1_AKT_NEG
PIK3C1_AKT_NEG, PIK3C1_AKT_POS, RAS_NEG
PIK3C1_AKT_NEG, PIK3C1_AKT_POS, RAS_NEG, RAS_POS, RB_POS, TP53_POS
PIK3C1_AKT_NEG, PIK3C1_AKT_POS, RAS_POS
PIK3C1_AKT_NEG, PIK3C1_AKT_POS, RAS_POS, RB_NEG, RB_POS, TP53_NEG, TP53_POS
PIK3C1_AKT_NEG, PIK3C1_AKT_POS, RAS_POS, RB_POS, TP53_POS
PIK3C1_AKT_NEG, RAS_NEG, RB_NEG
PIK3C1_AKT_NEG, RAS_POS
PIK3C1_AKT_NEG, RAS_POS, RB_NEG
PIK3C1_AKT_NEG, RAS_POS, RB_NEG, RB_POS, TP53_POS
PIK3C1_AKT_NEG, RAS_POS, RB_POS, TP53_NEG, TP53_POS
PIK3C1_AKT_NEG, RAS_POS, RB_POS, TP53_POS
PIK3C1_AKT_NEG, RB_NEG, TP53_NEG
PIK3C1_AKT_NEG, RB_POS
PIK3C1_AKT_NEG, RB_POS, TP53_POS
PIK3C1_AKT_NEG, TP53_NEG
PIK3C1_AKT_POS
PIK3C1_AKT_POS, RAS_NEG, RAS_POS, RB_NEG, RB_POS, TP53_POS
PIK3C1_AKT_POS, RAS_NEG, RAS_POS, RB_POS, TP53_POS
PIK3C1_AKT_POS, RAS_NEG, RB_POS, TP53_POS
PIK3C1_AKT_POS, RAS_POS
PIK3C1_AKT_POS, RAS_POS, RB_NEG
PIK3C1_AKT_POS, RAS_POS, RB_NEG, RB_POS, TP53_NEG, TP53_POS
PIK3C1_AKT_POS, RAS_POS, RB_NEG, RB_POS, TP53_POS
PIK3C1_AKT_POS, RAS_POS, RB_NEG, TP53_NEG
PIK3C1_AKT_POS, RAS_POS, RB_NEG, TP53_POS
PIK3C1_AKT_POS, RAS_POS, RB_POS, TP53_POS
PIK3C1_AKT_POS, RB_NEG
PIK3C1_AKT_POS, RB_NEG, TP53_NEG
PIK3C1_AKT_POS, RB_POS
PIK3C1_AKT_POS, RB_POS, TP53_POS
RAS_NEG
RAS_NEG, RAS_POS
RAS_NEG, RAS_POS, RB_POS, TP53_POS
RAS_NEG, RB_NEG
RAS_NEG, RB_POS, TP53_POS
RAS_POS
RAS_POS, RB_NEG
RAS_POS, RB_NEG, RB_POS
RAS_POS, RB_NEG, RB_POS, TP53_NEG, TP53_POS
RAS_POS, RB_NEG, RB_POS, TP53_POS
RAS_POS, RB_NEG, TP53_NEG
RAS_POS, RB_POS
RAS_POS, RB_POS, TP53_NEG, TP53_POS
RAS_POS, RB_POS, TP53_POS
RAS_POS, TP53_NEG
RAS_POS, TP53_POS
RB_NEG
RB_NEG, RB_POS
RB_NEG, RB_POS, TP53_POS
RB_NEG, TP53_NEG
RB_POS
RB_POS, TP53_POS
TP53_NEG
TP53_POS
WT
Jensen−Shannon distance
1.00
0.75
0.50
0.25
0.00
Comparison
TD vs. TD
CE vs. CE
CE vs. TD

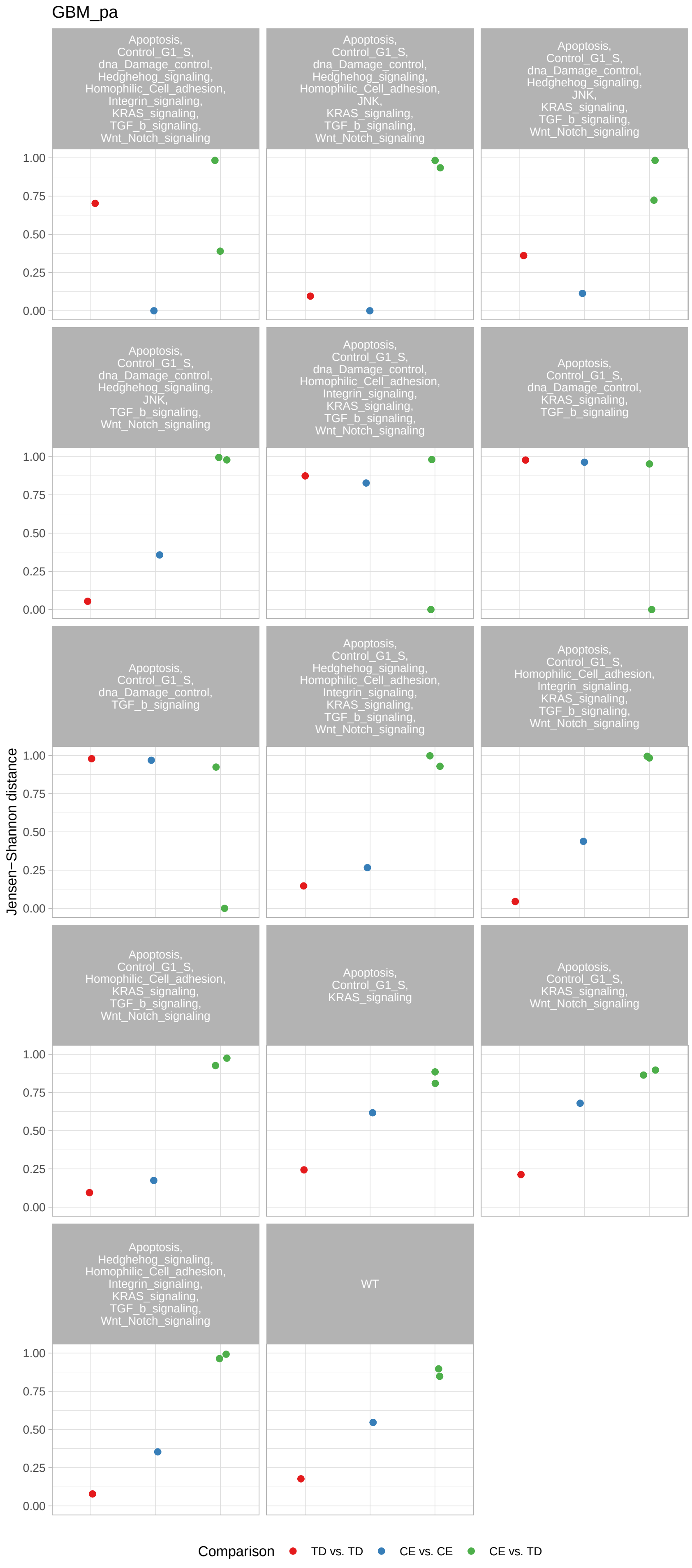
GBM_pa
Apoptosis, Control_G1_S, dna_Damage_control, Hedghehog_signaling, Homophilic_Cell_adhesion, Integrin_signaling, KRAS_signaling, TGF_b_signaling, Wnt_Notch_signaling
Apoptosis, Control_G1_S, dna_Damage_control, Hedghehog_signaling, Homophilic_Cell_adhesion, JNK, KRAS_signaling, TGF_b_signaling, Wnt_Notch_signaling
Apoptosis, Control_G1_S, dna_Damage_control, Hedghehog_signaling, JNK, KRAS_signaling, TGF_b_signaling, Wnt_Notch_signaling
Apoptosis, Control_G1_S, dna_Damage_control, Hedghehog_signaling, JNK, TGF_b_signaling, Wnt_Notch_signaling
Apoptosis, Control_G1_S, dna_Damage_control, Homophilic_Cell_adhesion, Integrin_signaling, KRAS_signaling, TGF_b_signaling, Wnt_Notch_signaling
Apoptosis, Control_G1_S, dna_Damage_control, KRAS_signaling, TGF_b_signaling
Apoptosis, Control_G1_S, dna_Damage_control, TGF_b_signaling
Apoptosis, Control_G1_S, Hedghehog_signaling, Homophilic_Cell_adhesion, Integrin_signaling, KRAS_signaling, TGF_b_signaling, Wnt_Notch_signaling
Apoptosis, Control_G1_S, Homophilic_Cell_adhesion, Integrin_signaling, KRAS_signaling, TGF_b_signaling, Wnt_Notch_signaling
Apoptosis, Control_G1_S, Homophilic_Cell_adhesion, KRAS_signaling, TGF_b_signaling, Wnt_Notch_signaling
Apoptosis, Control_G1_S, KRAS_signaling
Apoptosis, Control_G1_S, KRAS_signaling, Wnt_Notch_signaling
Apoptosis, Hedghehog_signaling, Homophilic_Cell_adhesion, Integrin_signaling, KRAS_signaling, TGF_b_signaling, Wnt_Notch_signaling
WT
Jensen−Shannon distance
1.00
0.75
0.50
0.25
0.00
Comparison
TD vs. TD
CE vs. CE
CE vs. TD

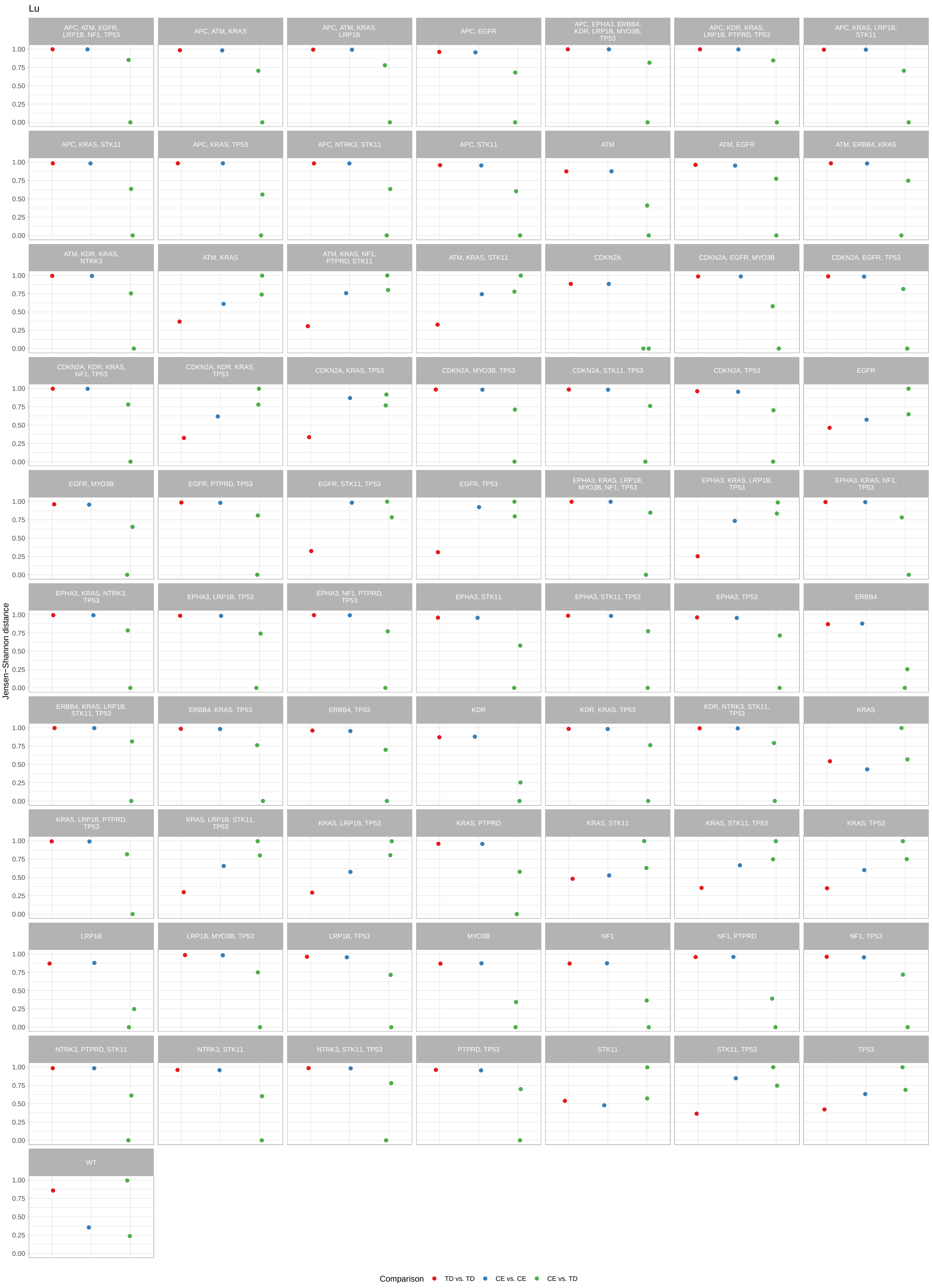
Lu
Jensen−Shannon distance
1.00
0.75
0.50
0.25
0.00
APC, ATM, EGFR, LRP1B, NF1, TP53
APC, ATM, KRAS
APC, ATM, KRAS, LRP1B
APC, EGFR
APC, EPHA3, ERBB4, KDR, LRP1B, MYO3B, TP53
APC, KDR, KRAS, LRP1B, PTPRD, TP53
APC, KRAS, LRP1B, STK11
APC, KRAS, STK11
APC, KRAS, TP53
APC, NTRK3, STK11
APC, STK11
ATM
ATM, EGFR
ATM, ERBB4, KRAS
ATM, KDR, KRAS, NTRK3
ATM, KRAS
ATM, KRAS, NF1, PTPRD, STK11
ATM, KRAS, STK11
CDKN2A
CDKN2A, EGFR, MYO3B
CDKN2A, EGFR, TP53
CDKN2A, KDR, KRAS, NF1, TP53
CDKN2A, KDR, KRAS, TP53
CDKN2A, KRAS, TP53
CDKN2A, MYO3B, TP53
CDKN2A, STK11, TP53
CDKN2A, TP53
EGFR
EGFR, MYO3B
EGFR, PTPRD, TP53
EGFR, STK11, TP53
EGFR, TP53
EPHA3, KRAS, LRP1B, MYO3B, NF1, TP53
EPHA3, KRAS, LRP1B, TP53
EPHA3, KRAS, NF1, TP53
EPHA3, KRAS, NTRK3, TP53
EPHA3, LRP1B, TP53
EPHA3, NF1, PTPRD, TP53
EPHA3, STK11
EPHA3, STK11, TP53
EPHA3, TP53
ERBB4
ERBB4, KRAS, LRP1B, STK11, TP53
ERBB4, KRAS, TP53
ERBB4, TP53
KDR
KDR, KRAS, TP53
KDR, NTRK3, STK11, TP53
KRAS
KRAS, LRP1B, PTPRD, TP53
KRAS, LRP1B, STK11, TP53
KRAS, LRP1B, TP53
KRAS, PTPRD
KRAS, STK11
KRAS, STK11, TP53
KRAS, TP53
LRP1B
LRP1B, MYO3B, TP53
LRP1B, TP53
MYO3B
NF1
NF1, PTPRD
NF1, TP53
NTRK3, PTPRD, STK11
NTRK3, STK11
NTRK3, STK11, TP53
PTPRD, TP53
STK11
STK11, TP53
TP53
WT
Comparison
TD vs. TD
CE vs. CE
CE vs. TD

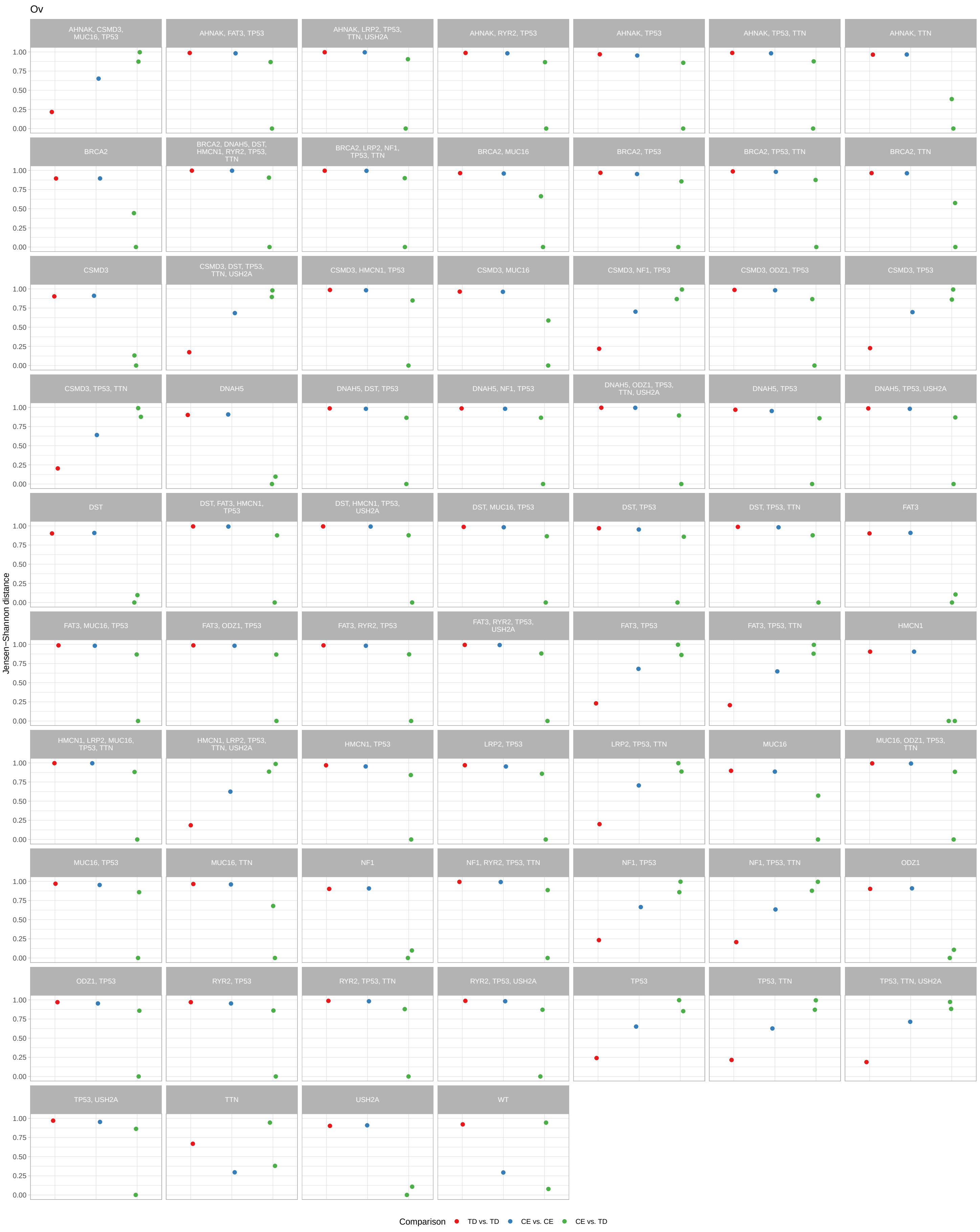

Ov
AHNAK, CSMD3, MUC16, TP53
AHNAK, FAT3, TP53
AHNAK, LRP2, TP53, TTN, USH2A
AHNAK, RYR2, TP53
AHNAK, TP53
AHNAK, TP53, TTN
AHNAK, TTN
BRCA2
BRCA2, DNAH5, DST, HMCN1, RYR2, TP53, TTN
BRCA2, LRP2, NF1, TP53, TTN
BRCA2, MUC16
BRCA2, TP53
BRCA2, TP53, TTN
BRCA2, TTN
CSMD3
CSMD3, DST, TP53, TTN, USH2A
CSMD3, HMCN1, TP53
CSMD3, MUC16
CSMD3, NF1, TP53
CSMD3, ODZ1, TP53
CSMD3, TP53
CSMD3, TP53, TTN
DNAH5
DNAH5, DST, TP53
DNAH5, NF1, TP53
DNAH5, ODZ1, TP53, TTN, USH2A
DNAH5, TP53
DNAH5, TP53, USH2A
DST
DST, FAT3, HMCN1, TP53
DST, HMCN1, TP53, USH2A
DST, MUC16, TP53
DST, TP53
DST, TP53, TTN
FAT3
FAT3, MUC16, TP53
FAT3, ODZ1, TP53
FAT3, RYR2, TP53
FAT3, RYR2, TP53, USH2A
FAT3, TP53
FAT3, TP53, TTN
HMCN1
HMCN1, LRP2, MUC16, TP53, TTN
HMCN1, LRP2, TP53, TTN, USH2A
HMCN1, TP53
LRP2, TP53
LRP2, TP53, TTN
MUC16
MUC16, ODZ1, TP53, TTN
MUC16, TP53
MUC16, TTN
NF1
NF1, RYR2, TP53, TTN
NF1, TP53
NF1, TP53, TTN
ODZ1
ODZ1, TP53
RYR2, TP53
RYR2, TP53, TTN
RYR2, TP53, USH2A
TP53
TP53, TTN
TP53, TTN, USH2A
TP53, USH2A
TTN
USH2A
WT
Jensen−Shannon distance
1.00
0.75
0.50
0.25
0.00
Comparison
TD vs. TD
CE vs. CE
CE vs. TD

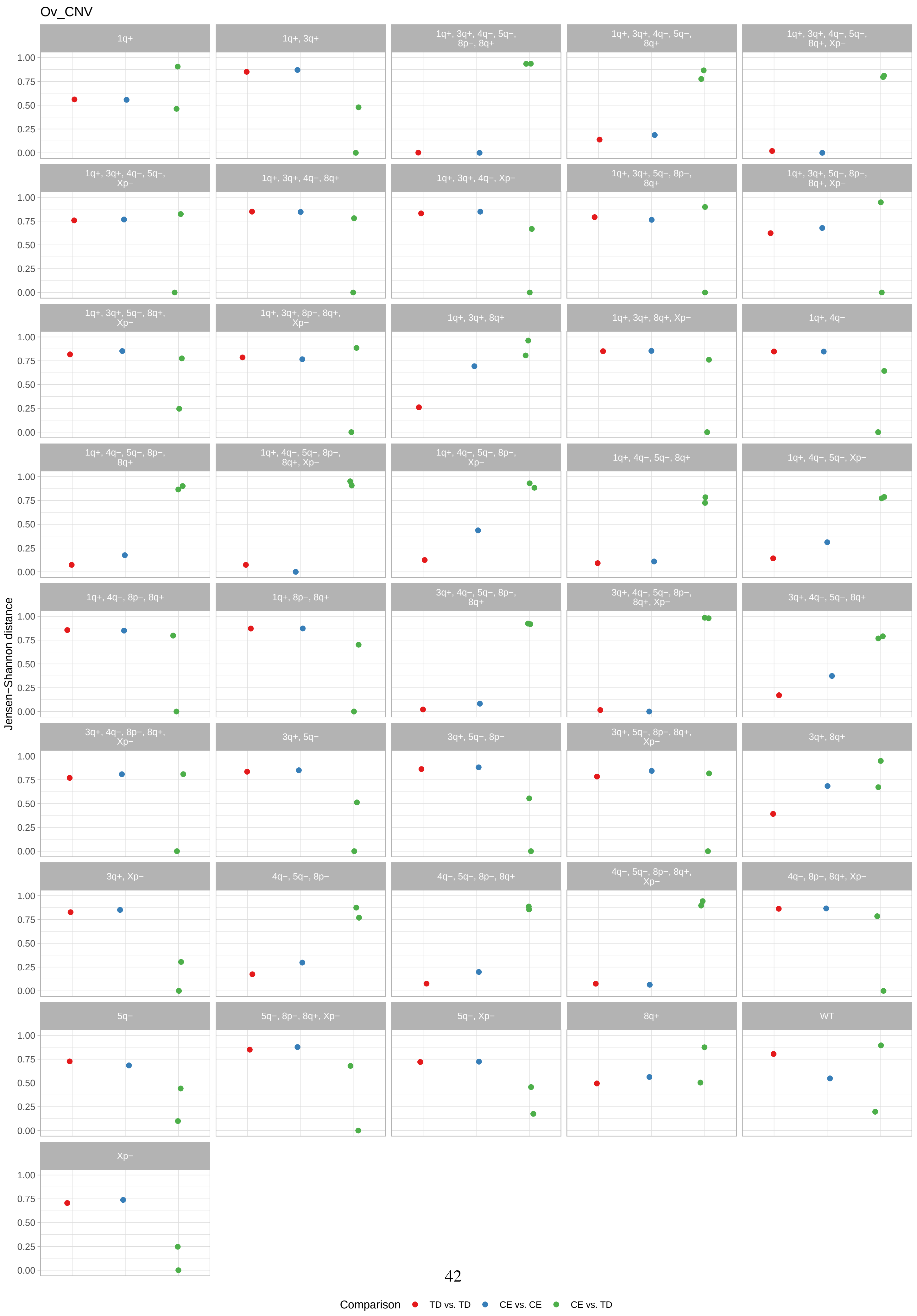

Ov_CNV
1q+
1q+, 3q+
1q+, 3q+, 4q−, 5q−, 8p−, 8q+
1q+, 3q+, 4q−, 5q−, 8q+
1q+, 3q+, 4q−, 5q−, 8q+, Xp−
1q+, 3q+, 4q−, 5q−, Xp−
1q+, 3q+, 4q−, 8q+
1q+, 3q+, 4q−, Xp−
1q+, 3q+, 5q−, 8p−, 8q+
1q+, 3q+, 5q−, 8p−, 8q+, Xp−
1q+, 3q+, 5q−, 8q+, Xp−
1q+, 3q+, 8p−, 8q+, Xp−
1q+, 3q+, 8q+
1q+, 3q+, 8q+, Xp−
1q+, 4q−
1q+, 4q−, 5q−, 8p−, 8q+
1q+, 4q−, 5q−, 8p−, 8q+, Xp−
1q+, 4q−, 5q−, 8p−, Xp−
1q+, 4q−, 5q−, 8q+
1q+, 4q−, 5q−, Xp−
1q+, 4q−, 8p−, 8q+
1q+, 8p−, 8q+
3q+, 4q−, 5q−, 8p−, 8q+
3q+, 4q−, 5q−, 8p−, 8q+, Xp−
3q+, 4q−, 5q−, 8q+
3q+, 4q−, 8p−, 8q+, Xp−
3q+, 5q−
3q+, 5q−, 8p−
3q+, 5q−, 8p−, 8q+, Xp−
3q+, 8q+
3q+, Xp−
4q−, 5q−, 8p−
4q−, 5q−, 8p−, 8q+
4q−, 5q−, 8p−, 8q+, Xp−
4q−, 8p−, 8q+, Xp−
5q−
5q−, 8p−, 8q+, Xp−
5q−, Xp−
8q+
WT
Xp−
Jensen−Shannon distance
0.00
0.25
0.50
0.75
1.00
Comparison
TD vs. TD
CE vs. CE
CE vs. TD

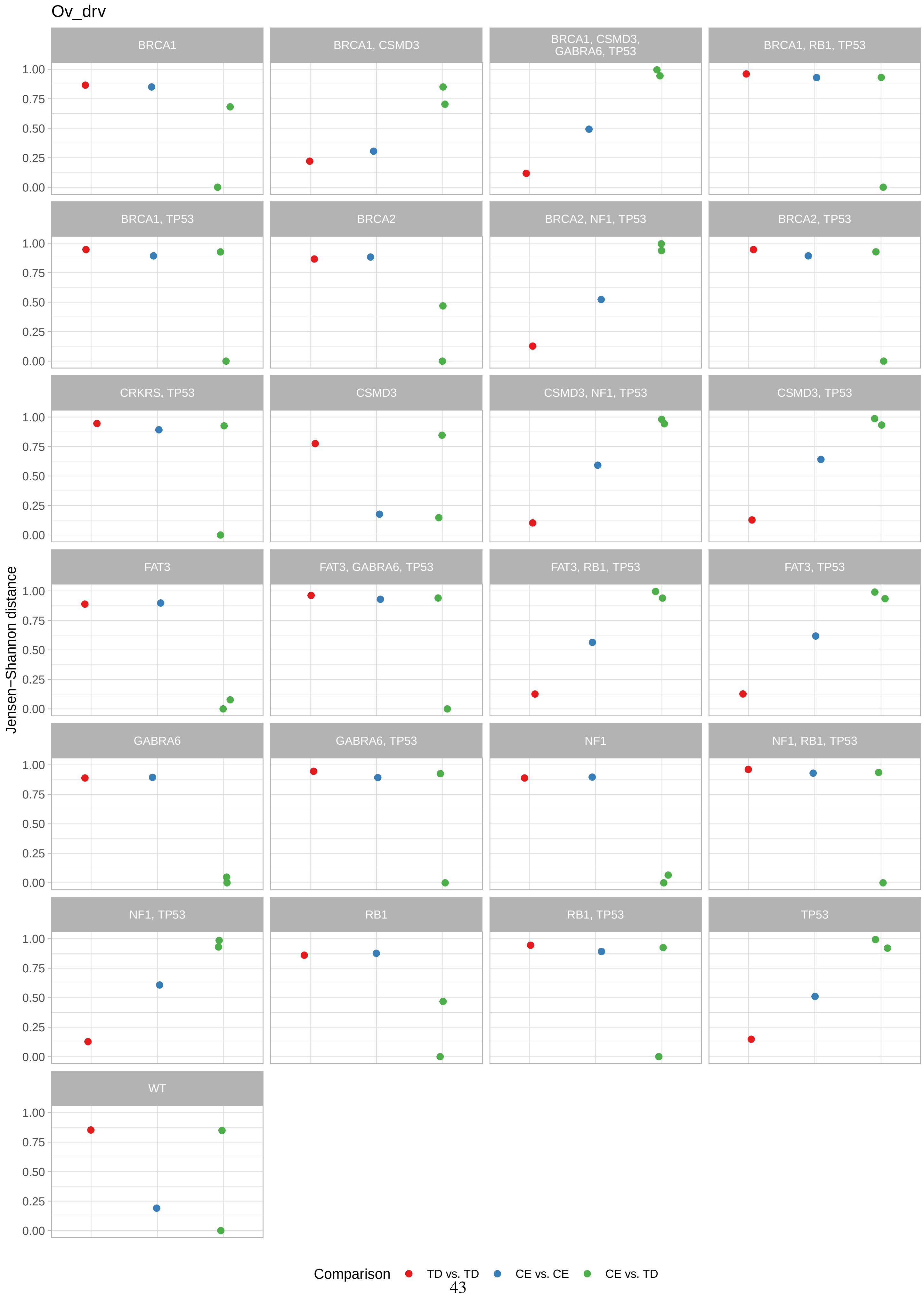

Ov_drv
BRCA1
BRCA1, CSMD3
BRCA1, CSMD3, GABRA6, TP53
BRCA1, RB1, TP53
BRCA1, TP53
BRCA2
BRCA2, NF1, TP53
BRCA2, TP53
CRKRS, TP53
CSMD3
CSMD3, NF1, TP53
CSMD3, TP53
FAT3
FAT3, GABRA6, TP53
FAT3, RB1, TP53
FAT3, TP53
GABRA6
GABRA6, TP53
NF1
NF1, RB1, TP53
NF1, TP53
RB1
RB1, TP53
TP53
WT
Jensen−Shannon distance
1.00
0.75
0.50
0.25
0.00
Comparison
TD vs. TD
CE vs. CE
CE vs. TD

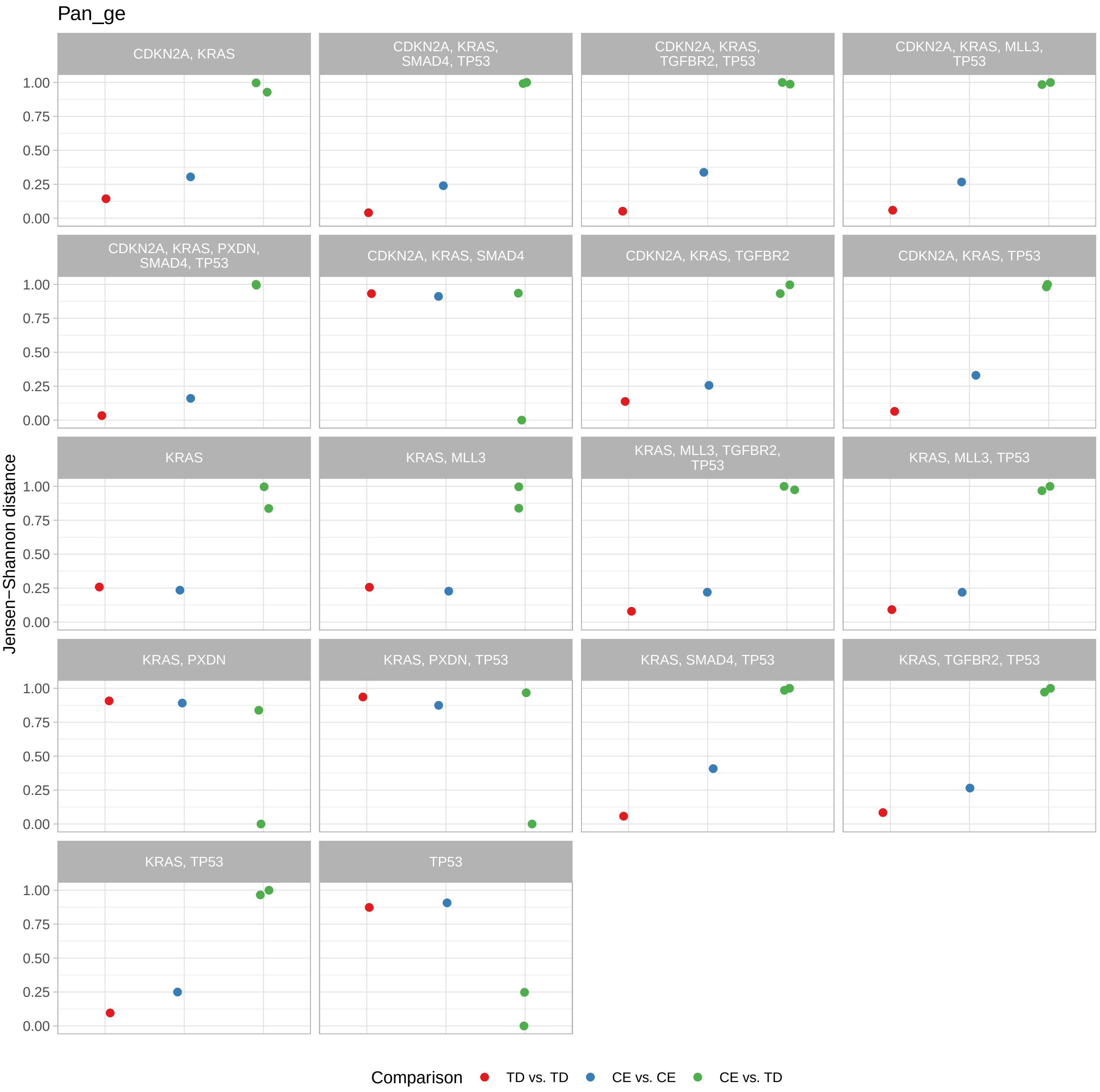

Pan_ge
CDKN2A, KRAS
CDKN2A, KRAS, SMAD4, TP53
CDKN2A, KRAS, TGFBR2, TP53
CDKN2A, KRAS, MLL3, TP53
CDKN2A, KRAS, PXDN, SMAD4, TP53
CDKN2A, KRAS, SMAD4
CDKN2A, KRAS, TGFBR2
CDKN2A, KRAS, TP53
KRAS
KRAS, MLL3
KRAS, MLL3, TGFBR2, TP53
KRAS, MLL3, TP53
KRAS, PXDN
KRAS, PXDN, TP53
KRAS, SMAD4, TP53
KRAS, TGFBR2, TP53
KRAS, TP53
TP53
Jensen−Shannon distance
1.00
0.75
0.50
0.25
0.00
Comparison
TD vs. TD
CE vs. CE
CE vs. TD

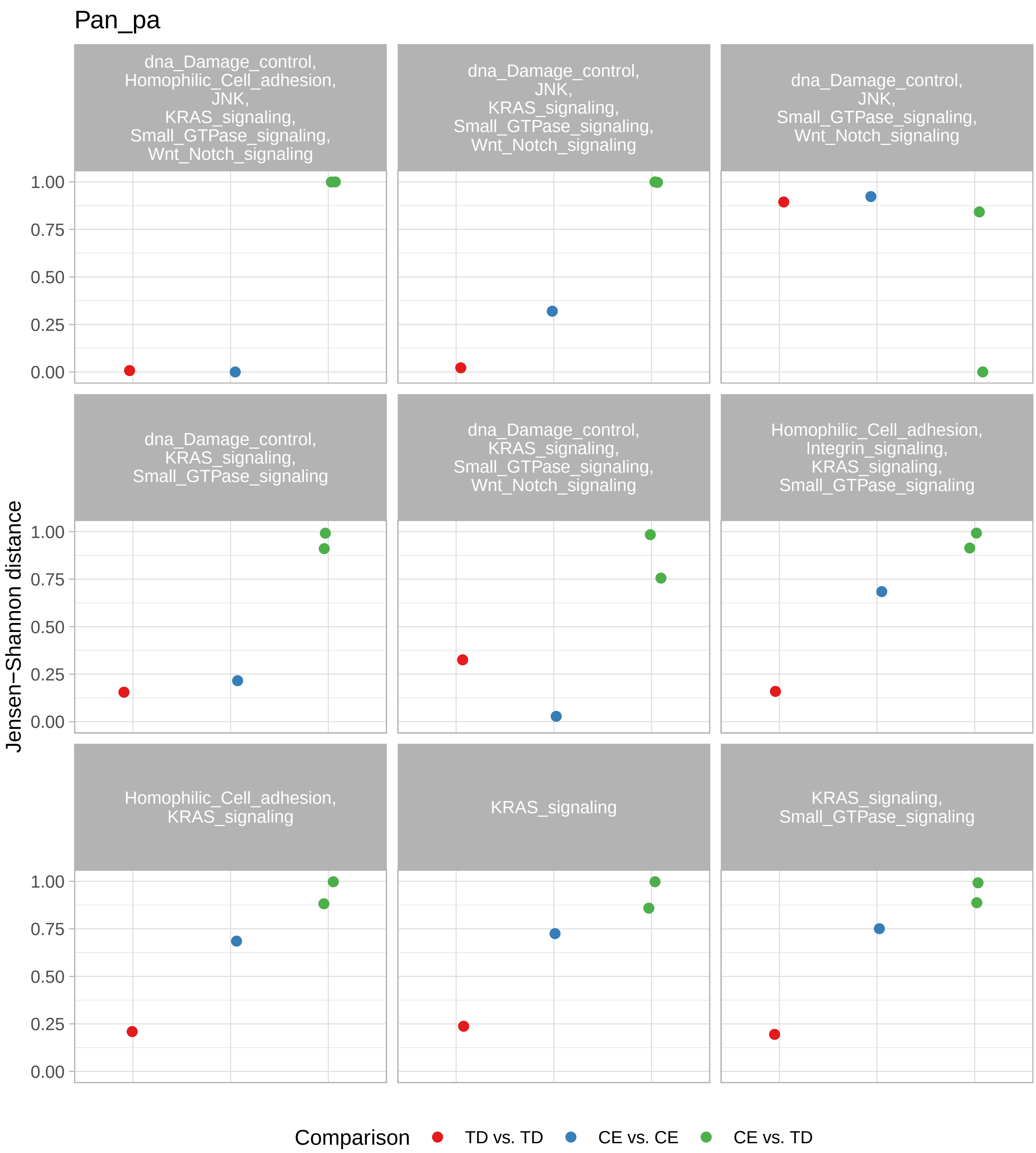

Pan_pa
dna_Damage_control, Homophilic_Cell_adhesion, JNK, KRAS_signaling, Small_GTPase_signaling, Wnt_Notch_signaling
dna_Damage_control, JNK, KRAS_signaling, Small_GTPase_signaling, Wnt_Notch_signaling
dna_Damage_control, JNK, Small_GTPase_signaling, Wnt_Notch_signaling
dna_Damage_control, KRAS_signaling, Small_GTPase_signaling
dna_Damage_control, KRAS_signaling, Small_GTPase_signaling, Wnt_Notch_signaling
Homophilic_Cell_adhesion, Integrin_signaling, KRAS_signaling, Small_GTPase_signaling
Homophilic_Cell_adhesion, KRAS_signaling
KRAS_signaling
KRAS_signaling, Small_GTPase_signaling
Jensen−Shannon distance
1.00
0.75
0.50
0.25
0.00
Comparison
TD vs. TD
CE vs. CE
CE vs. TD

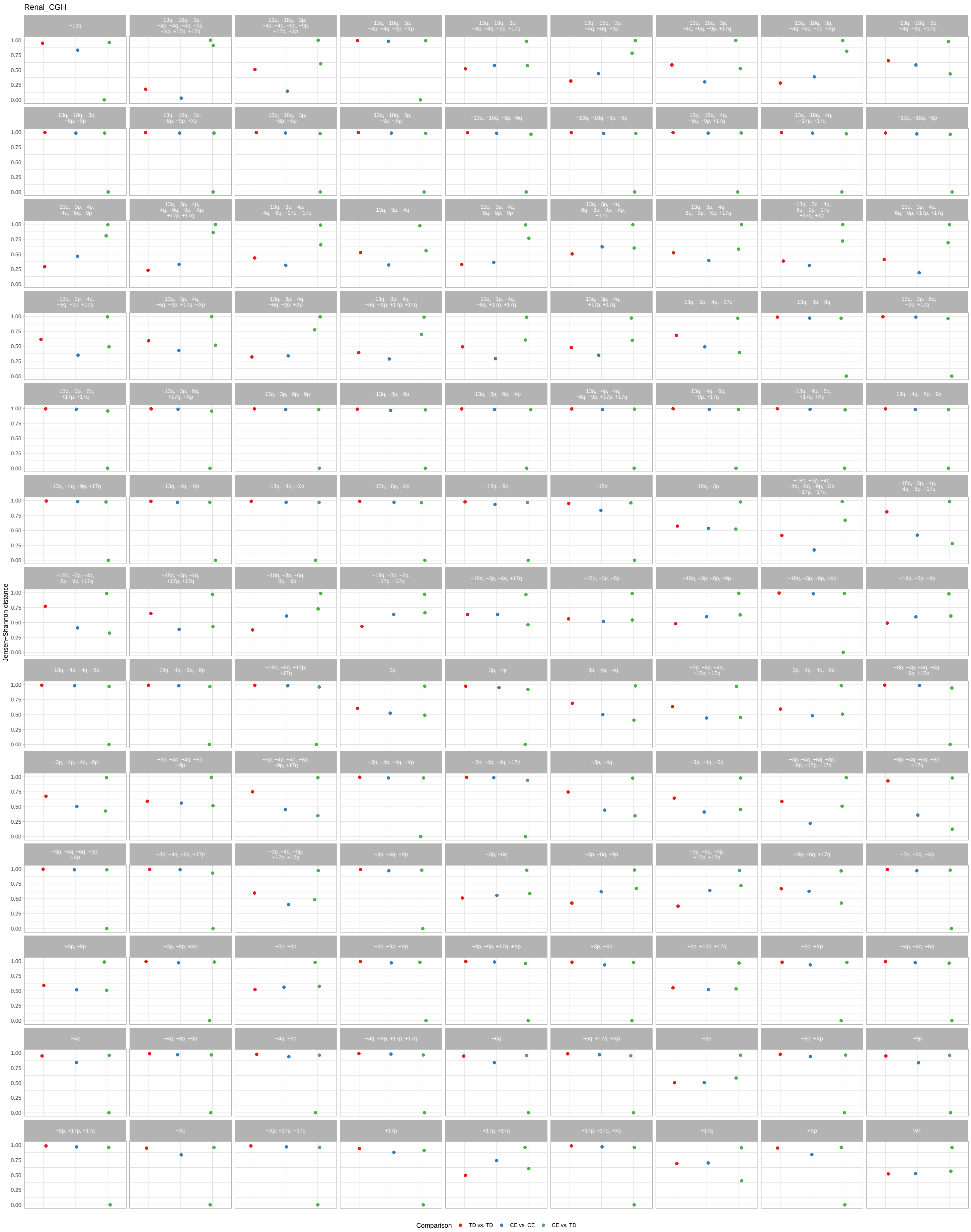
Renal_CGH
Jensen−Shannon distance
0.00
0.25
0.50
0.75
1.00
−13q
−13q, −18q, −3p, −4p, −4q, −6q, −9p, −Xp, +17p, +17q
−13q, −18q, −3p, −4p, −4q, −6q, −9p, +17q, +Xp
−13q, −18q, −3p, −4p, −4q, −9p, −Xp
−13q, −18q, −3p, −4p, −4q, −9p, +17q
−13q, −18q, −3p, −4q, −6q, −9p
−13q, −18q, −3p, −4q, −6q, −9p, +17q
−13q, −18q, −3p, −4q, −6q, −9p, +Xp
−13q, −18q, −3p, −4q, −6q, +17q
−13q, −18q, −3p, −8p, −9p
−13q, −18q, −3p, −8p, −9p, +Xp
−13q, −18q, −3p, −8p, −Xp
−13q, −18q, −3p, −9p, −Xp
−13q, −18q, −3p, −6q
−13q, −18q, −3p, −9p
−13q, −18q, −4q, −6q, −9p, +17q
−13q, −18q, −4q, +17p, +17q
−13q, −18q, −8p
−13q, −3p, −4p, −4q, −6q, −9p
−13q, −3p, −4p, −4q, −6q, −9p, −Xp, +17p, +17q
−13q, −3p, −4p, −4q, −6q, +17p, +17q
−13q, −3p, −4q
−13q, −3p, −4q, −6q, −8p, −9p
−13q, −3p, −4q, −6q, −8p, −9p, −Xp, +17q
−13q, −3p, −4q, −6q, −9p, −Xp, +17q
−13q, −3p, −4q, −6q, −9p, +17p, +17q, +Xp
−13q, −3p, −4q, −6q, −9p, +17p, +17q
−13q, −3p, −4q, −6q, −9p, +17q
−13q, −3p, −4q, −6q, −9p, +17q, +Xp
−13q, −3p, −4q, −6q, −9p, +Xp
−13q, −3p, −4q, −6q, −Xp, +17p, +17q
−13q, −3p, −4q, −6q, +17p, +17q
−13q, −3p, −4q, +17p, +17q
−13q, −3p, −4q, +17q
−13q, −3p, −6q
−13q, −3p, −6q, −9p, +17q
−13q, −3p, −6q, +17p, +17q
−13q, −3p, −6q, +17q, +Xp
−13q, −3p, −8p, −9p
−13q, −3p, −9p
−13q, −3p, −9p, −Xp
−13q, −4p, −4q, −6q, −9p, +17p, +17q
−13q, −4q, −6q, −9p, +17q
−13q, −4q, −6q, +17q, +Xp
−13q, −4q, −8p, −9p
−13q, −4q, −9p, +17q
−13q, −4q, −Xp
−13q, −4q, +Xp
−13q, −8p, −Xp
−13q, −9p
−18q
−18q, −3p
−18q, −3p, −4p, −4q, −6q, −9p, −Xp, +17p, +17q
−18q, −3p, −4q, −8p, +17q
−18q, −3p, −4q, −8p, −9p, +17q
−18q, −3p, −4q, +17p, +17q
−18q, −3p, −6q, −8p, −9p
−18q, −3p, −6q, +17p, +17q
−18q, −3p, −6q, +17q
−18q, −3p, −8p
−18q, −3p, −8p, −9p
−18q, −3p, −8p, −Xp
−18q, −3p, −9p
−18q, −4p, −4q, −8p
−18q, −4q, −6q, −9p
−18q, −6q, +17p, +17q
−3p
−3p, −4p
−3p, −4p, −4q
−3p, −4p, −4q, +17p, +17q
−3p, −4p, −4q, −6q
−3p, −4p, −4q, −6q, −9p, +17p
−3p, −4p, −4q, −8p
−3p, −4p, −4q, −8p, −9p
−3p, −4p, −4q, −8p, −9p, +17q
−3p, −4p, −4q, −Xp
−3p, −4p, −4q, +17p
−3p, −4q
−3p, −4q, −6q
−3p, −4q, −6q, −8p, −9p, +17p, +17q
−3p, −4q, −6q, −8p, +17q
−3p, −4q, −6q, −9p, +Xp
−3p, −4q, −6q, +17p
−3p, −4q, −9p, +17p, +17q
−3p, −4q, −Xp
−3p, −6q
−3p, −6q, −9p
−3p, −6q, −9p, +17p, +17q
−3p, −6q, +17q
−3p, −6q, +Xp
−3p, −8p
−3p, −8p, +Xp
−3p, −9p
−3p, −9p, −Xp
−3p, −9p, +17q, +Xp
−3p, −Xp
−3p, +17p, +17q
−3p, +Xp
−4p, −4q, −6q
−4q
−4q, −8p, −9p
−4q, −9p
−4q, −Xp, +17p, +17q
−6q
−6q, +17q, +Xp
−8p
−8p, +Xp
−9p
−9p, +17p, +17q
−Xp
−Xp, +17p, +17q
+17p
+17p, +17q
+17p, +17q, +Xp
+17q
+Xp
WT
Comparison
TD vs. TD
CE vs. CE
CE vs. TD
